# Feedback between PI4P signaling and ER-PM contact sites orchestrates polarized root hair growth

**DOI:** 10.64898/2026.06.26.734759

**Authors:** Vedrana Marković, Vincent Bayle, Gwennogan Dubois, Frédérique Rozier, Vítor Amorim-Silva, Jorge Morello-López, Sacha Grenet, Selene García-Hernández, Miguel A. Botella, Yvon Jaillais

## Abstract

Eukaryotic cells are composed of different organelles that communicate with one another through direct contacts, which are necessary for a host of cellular reactions and for responding to different developmental and environmental changes. Plasma membrane (PM) forms extensive contacts with the endoplasmic reticulum (ER) at specific sites named ER-PM contact sites. These contacts play crucial functions in lipid homeostasis, Ca^2+^ regulation and signaling in all eukaryotes. However, the mechanisms by which plant ER-PM contact site proteins tether to the PM, as well as the dynamics of these contact sites, remain poorly understood. Here, we investigate the importance of phosphoinositides in the establishment and dynamics of ER-PM contact site proteins in plants. We found that phosphatidylinositol-4-phosphate (PI4P), rather than phosphatidylinositol 4,5-bisphosphate (PI(4,5)P_2_), is required for the association of ER-PM contact site proteins with the PM. Furthermore, we identified a PI4P phosphatase, SUPPRESSOR-OF-ACTIN7 (SAC7), that associates with the ER-PM contact site protein SYNAPTOTAGMIN1 (SYT1) and regulates its dynamic association with the PM. In particular, we found that in growing root hairs, a highly polarized cell type, SAC7 removes SYT1-containing contact sites at the growing tip. Consistently, optogenetic induction of ER–PM tethering reduced root hair elongation within minutes of blue light induction. Altogether, we propose a link between SAC7-mediated regulation of PI4P, dynamic ER-PM contact site establishment and polarized cell growth in plants.

## Introduction

Eukaryotic cells are characterized by the presence of membrane-bound organelles that compartmentalize essential cellular functions. Increasing evidence indicates that these organelles are not isolated structures, but instead form dynamic and interconnected networks through membrane contact sites, enabling the exchange of signals, lipids, metabolites, and ions^1,2^. Membrane contact sites are specialized regions where the membranes of two distinct organelles are held in close proximity—typically within 10–30 nanometers—without fusing. They are widely present in all eukaryotic cells and for the most part they include the interaction of the Endoplasmic Reticulum (ER) membrane with another cellular membrane, such as PM^3^. It was shown that the ER-PM contact sites play role in many processes crucial for cellular development including lipid transfer between membranes^4–6^, receptor kinase signaling^7^, autophagosome formation^8^, and calcium homeostasis^9–11^.

The close proximity between organelles at membrane contact sites is stabilized by tethering proteins that physically bridge adjacent organelles. Tethering proteins or tethering complexes that are found at the ER-PM contact sites contain transmembrane region embedded in the ER membrane and domains that associate with the PM. To interact with the PM, these proteins utilize specific lipid-binding domains that interact with anionic lipids present at the PM^3^. In yeast and animal cells, presence of specific anionic lipids at the PM is crucial for the formation of the ER-PM contact sites^12,13^.

Several families of ER–PM tethering proteins have been identified in plants^14^. Among them, SYNAPTOTAGMIN1 (SYT1), a member of the plant synaptotagmin (SYT) family and an ortholog of animal extended synaptotagmins (E-SYTs) and yeast tricalbins, is one of the best-characterized ER–PM tethers in plants^15,16^. SYT1 functions as a lipid transfer protein that removes diacylglycerol (DAG) from the PM and transfers it to the ER, thereby preserving PM integrity during abiotic stress^17,18^. Localization of SYT1 at ER–PM contact sites is mediated by an ER-anchoring transmembrane domain and two C2 domains that bind anionic lipids at the PM^15,19^.

Another class of ER–PM tethers is represented by the VAP27 family, orthologs of the yeast Scs2 proteins^20,21^. VAP27 is an ER-resident protein containing a major sperm protein (MSP) domain that mediates interactions with members of the NETWORKED (NET) family of actin-binding proteins, which link the actin cytoskeleton to endomembrane compartments. Among these, NETWORKED 3C (NET3C) interacts with VAP27 and connects the complex to the PM. Thus, the VAP27–NET3C complex forms a molecular bridge between the ER and PM, physically coupling the two membranes to the actin cytoskeleton^22^.

Despite these advances, the lipid determinants underlying the formation and specialization of distinct plant ER–PM contact sites remain a matter of debate. In particular, whether specific lipid environments contribute to the differential localization and dynamics of individual tethering complexes is unknown, as is how these mechanisms impact plant development and physiology.

Addressing the question of which lipids target which tether to the PM is difficult for several reasons. First, interaction data obtained *in vitro* may not reflect the actual situation *in vivo* since lipid binding is often promiscuous^23^. Second, phosphoinositide species are highly interdependent^21,24^. This means that interfering with one phosphoinositide species may impact the level of another. This difficulty in studying tether-lipid interaction is best illustrated with the Arabidopsis SYT1 protein. Using protein-lipid overlay assays and liposome sedimentation, SYT1 was shown to bind to several anionic phospholipids, including phosphatidylserine (PS) in a calcium dependent manner, as well as phosphoinositides including PI4P and PI(4,5)P_2_^15,19,25^. Inspired by studies on animal E-SYTs^12^, several detailed characterizations of SYT1-PI(4,5)P_2_ interaction were carried out *in vitro*^19^. These studies showed that PI(4,5)P_2_ can recapitulate the ability of SYT1 to tether liposomes *in vitro*. This work falls perfectly in line with previous observations suggesting that the amount of SYT1-mediated contact sites correlates with an increase in PI(4,5)P_2_ at the PM during salt stress^17^. Together, these data suggest that like in animal cells, PI(4,5)P_2_ could be the main lipid that regulates the formation of SYT1-mediated ER-PM contacts, although *in vivo* proof of this is lacking. At the same time, we should also take into consideration that PI(4,5)P_2_ is present at very low concentrations in the PM of plant cells, while PI4P is the main phosphoinositide species present in this compartment^26–28^. Accordingly, removal of PI4P at the PM detaches SYT1 from the PM^6^. This data suggests an important role for PI4P in SYT1 attachment to the PM, but we cannot rule out that removal of PI4P also impacts the levels of PI(4,5)P_2_.

Here, we set out to test whether PI4P, PI(4,5)P_2_ or both lipids species are important for the establishment of ER-PM contacts in *Nicotiana benthamiana* leaves and *Arabidopsis thaliana* root cells. To this end, we used a series of genetic tools that allow the transient depletion of PI4P or PI(4,5)P_2_ in plant cells. Using this approach, we concluded that PI4P, not PI(4,5)P_2,_ is the main phosphoinositide for ER-PM targeting, at least in basal —unstressed—conditions. Moreover, using an interactomic approach, we found that SYT1 forms a complex with a PI4P 4-phosphatase called SUPPRESSOR OF ACTIN7 (SAC7). SAC7 is an ER-localized transmembrane protein involved in PI4P signaling, cell-cell communication and root hair growth^29–32^. High spatiotemporal live imaging experiments suggested that SAC7 and SYT1 act in a feedback loop for reciprocal regulation of their localization.

We found that this regulatory mechanism is particularly important during root hair growth, where SAC7 negatively regulates the formation of stable ER–PM contacts at the growing tip. Consistently, optogenetic induction of ER–PM tethering reduced root hair elongation within minutes. Together, these findings identify SAC7 as a key regulator linking PI4P signaling with ER–PM contact site formation and dynamics in plants, with direct consequences on cell morphogenesis.

## Results

### PI(4,5)P_2_ is not required for the formation of ER–PM contact sites in *Nicotiana benthamiana* leaves

In animals, the presence of PI(4,5)P_2_ at the PM plays a crucial role in the establishment of ER-PM contact sites^12^, while PI4P is important for the lipid exchanges between the PM and the ER membrane^4,33,34^. Previous studies suggest that PI(4,5)P_2_ may also be involved in the PM localization of Arabidopsis SYT1^17^, while the tethering mechanism of other plant ER-PM contact site proteins remains unknown. To understand how ER-PM contact site proteins target the PM *in planta*, we investigated two well-characterized ER-PM contact site proteins in plants — SYT1 and NET3C. To test the importance of PI(4,5)P_2_ in the PM localization of SYT1 and NET3C *in planta*, we used the iDePP system, which is based on catalytically active or inactive (dead) *Drosophila* 5-phosphatase OCRL domain (*dm*OCRL) artificially targeted to the PM with a MAP (myristoylation and palmitoylation) sequence and tagged with mCherry for visualization^35^ (**Fig.1a**). We used transient expression in *Nicotiana benthamiana* leaf epidermis, a cell type with stable SYT1 and NET3C-mediated ER-PM contacts^22,36,37^ (**Supplementary Fig.1a**). In control conditions, when expressed alone or co-expressed with the MAP-*dm*OCRLdead, both SYT1-GFP and NET3C-GFP localized at ER-PM contact sites (**Figs.1b-c, left and middle panels**). Upon depletion of PI(4,5)P_2_ from the PM, by co-expression with MAP-dmOCRL, the localization of SYT1-GFP and NET3C-GFP at ER-PM contact sites was not altered (**Figs.1b-c, right panels**). To quantify the density of contact sites labeled by SYT1-GFP and NET3C-GFP, we automatically segmented them using the machine learning-based software ilastik^38^ and focusing on the round dotted structures^18^. Indeed, these dots correspond to the persistent signal observed during time lapses, which is a hallmark of ER-PM contact sites in plants^39–41^ (**Supplementary Figs. 1a-b**). The quantification of contact site density did not show a significant difference when SYT1-GFP and NET3C-GFP were co-expressed with either MAP–*dm*OCRL and MAP–*dm*OCRLdead (**Figs.1d-e**). Thus, PI(4,5)P_2_ is dispensable for the establishment of ER-PM contact sites in *Nicotiana bentamiana* leaves.

### PI4P dephosphorylation dissociates SYT1 and NET3C from ER-PM contact sites in *Nicotiana benthamiana* leaves

Next, we employed a similar strategy to deplete PI4P from the PM. We previously established a system that consists on the fusion of the active or inactive (dead) phosphatase domain of the yeast PI4P-phosphatase SAC1 (SAC1), with the MAP sequence, that induces PM targeting, and mCherry fluorescent protein for visualization^26^. Our previous experience with the iDePP system showed that yeast or mammal enzymes may not be fully active when expressed in plant cells because 20°C is a suboptimal temperature for their activities. We thus decided to improve our system by using the catalytic domain of *Drosophila melanogaster* SAC1 (*dm*SAC1) instead of the *Saccharomyces cerivisiae* SAC1 domain (**Fig.2a**). We verified the activity of these new constructs by co-expressing them with the PI4P biosensor mCitrine-1xPH^FAPP1^. In plant cells, PI4P mainly accumulates at the PM, but is also present, albeit at a much lower amount, in the TGN/endosomes. Thus, in unperturbed cells, PI4P biosensors localize at the PM, and depending on the biosensor, not or very weakly at the TGN/endosomes^26,28^. However, upon depletion of PI4P at the PM, the biosensors are free to bind the second PI4P pool at the TGN/endosomes and re-localize to these compartments. We found that the expression of MAP-*dm*SAC1, but not MAP–*dm*SAC1dead, caused a strong and reliable re-localization of this PI4P biosensor to TGN/endosomes (**Supplementary Figs. 2a-c**). This result confirmed that our system efficiently reduces PI4P amount at the PM.

Next, we used this system to investigate the localization of SYT1 and NET3C upon PI4P PM depletion. We co-expressed MAP-*dm*SAC1 or MAP–*dm*SAC1dead with the GFP-tagged variant of SYT1 and NET3C. In control conditions, when SYT1 and NET3C were expressed alone or with MAP–*dm*SAC1dead, both SYT1-GFP and NET3C-GFP localized in ER-PM contact sites in cortical regions of *Nicotiana benthamiana* leaf epidermal cells (**Figs.2b-c, left and middle panels**). However, when expressed with MAP-*dm*SAC1, the localization of SYT1-GFP and NET3C-GFP was altered (**Figs.2b-c, right panels**). Quantitative analysis of SYT1 and NET3C contact site density showed significant decrease of contact sites labeled by SYT1 and NET3C upon PI4P depletion (**Figs.2d-e**). Instead of its classical localization at ER-PM contact sites characterized by a cortical beads-on-strings pattern, upon PI4P depletion SYT1-GFP was organized as a cortical network reminiscent of the bulk ER, as previously observed^6,18^ (**Fig.2b, right panel**). We therefore transiently co-expressed MAP–*dm*SAC1 or MAP–*dm*SAC1dead with both SYT1-GFP and HDEL-CFP. HDEL-CFP is secreted in the lumen of the ER and as such is largely depleted from ER-PM contact sites^20^. Indeed, co-expression with the catalytically inactive MAP–*dm*SAC1dead showed that the dot-like structures labeled by SYT1-GFP were apposed to the tubular and sheet-like structures labeled by the ER luminal marker HDEL-CFP (**Fig.2f**). By contrast, co-expression with the active MAP–*dm*SAC1 enzyme led to a very strong overlap between SYT1-GFP and HDEL-CFP within ER tubules and the concomitant disappearance of SYT1-GFP signal within the stable dot-like structures (**Fig.2g**), indicating SYT1 re-localization from ER-PM contact sites into the ER upon PI4P depletion.

Upon PI4P depletion by MAP–*dm*SAC1, NET3C-GFP labeling in dot-like structures disappeared and instead localized to long fibers spanning the entire cell (**Fig.2c, right panel**). Because NET3C is known to bridge ER-PM contact sites with the actin cytoskeleton, we reasoned that NET3C might relocalize to actin fibers upon PI4P depletion. We thus transiently co-expressed MAP–*dm*SAC1 or MAP–*dm*SAC1dead with both NET3C-GFP and the actin marker Lifeact-mTURQUOISE2 (hereafter Lifeact-mTU2). This analysis confirmed that NET3C-GFP-labeled contact sites were closely positioned to actin cables when PI4P levels were unperturbed (co-expression with MAP–*dm*SAC1dead) ^22^ (**Fig.2h**), but disappeared from contact sites upon PI4P depletion and colocalized instead with the actin marker (**Fig.2i**). We also noticed that NET3C-GFP localized to big rounded structures upon co-expression with MAP–*dm*SAC1 (**Fig.2c, right panel**). These structures did not colocalize with Lifeact-mTU2 (**Fig.2i**), and given their number and size might represent an association with the chloroplast envelop.

Altogether, our results suggest that removal of PI4P, but not PI(4,5)P_2_, strongly impacts the localization of both SYT1 and NET3C in ER-PM contact sites in *Nicotiana benthamiana* leaves in basal conditions. Moreover, upon PI4P depletion at the PM, SYT1 relocalizes to the entire ER network, likely by diffusing within the ER membranes and NET3C associates with actin and possibly the chloroplast envelops.

### SYT1-GFP rapidly dissociates from contact sites upon PI4P depletion at the PM

Our results showed that PI4P depletion at the PM triggers the dissociation of both SYT1-GFP and NET3C-GFP from ER-PM contact sites in plants, and thus suggest a key role for this lipid in the formation of these structures. However, in our experiments, we observed *Nicotiana benthamiana* leaves 48 hours after infiltration, allowing time for lipid compensation. In particular, PI4P is a precursor to PI(4,5)P₂, and its depletion might affect PI(4,5)P₂ levels. To test if detachment of ER-PM contact site proteins from the PM occurs specifically as a response to the PI4P loss, we used a rapamycin-based system to target the catalytic domain of the SAC1 phosphatase to the PM in an induced and rapid manner^42^. This system employs the rapamycin-induced heterodimerization between the FKBP domain fused with yeast SAC1 and mCherry tag (FKBP-mCherry-SAC1) and FRB domain fused with MAP sequence and BFP2 tag (MAP-BFP2-FRB).

Therefore, upon rapamycin treatment FKBP-mCherry-SAC1 is recruited to the PM via interaction with MAP-BFP2-FRB that acts as a PM anchor (**Fig. 3a**). To test if this system can deplete PI4P *in planta*, Winkler *et al.* co-expressed it with a PI4P biosensor in *Nicotiana benthamiana* leaf epidermal cells, and showed that PI4P biosensor is delocalized from the PM to endosomes only when they used SAC1 but not the SAC1dead^42^. To investigate the effects of acute PI4P depletion on ER-PM contact site localization, we transiently co-expressed this system with SYT1-GFP in *Nicotiana benthamiana* leaf epidermal cells. Without rapamycin treatment, SAC1 and SAC1dead phosphatases were not expressed at the PM and SYT1 localized in contact sites (**Fig.3b and 3d**). Rapamycin addition targeted both SAC1 and SAC1dead to the PM within minutes (**Figs.3c and 3e, middle panels**). While a precise timing was difficult to record, the recruitment to the PM was as rapid as the time needed to inject the rapamycin and observe the sample. Concomitant with this rapid relocalization of FKBP-mCherry-SAC1 to the PM, we observed an immediate shift in SYT1-GFP localization from the contact sites to the bulk ER (**Fig.3c, left panel**), but not when the inactive enzyme was used (**Fig.3e, left panel**). Quantification analysis showed a significant decrease in the density of SYT1-GFP contact sites only upon rapamycin-inducible SAC1 recruitment to the PM, but not SAC1dead. Furthermore, rapamycin treatment alone did not change the density of SYT1-GFP-labeled contact sites in comparison to the control (**Fig.3f**). Dynamic analyses showed that SYT1-GFP-containing contact sites remained static within 1-minute time lapses when SAC1dead was recruited to the PM upon rapamycin treatment (**Fig.3g**). In contrast, SYT1-GFP localized in highly dynamic ER tubules when SAC1 was recruited to the PM, as revealed by persistency mapping (**Fig.3h**). These results show that the rapid depletion of PI4P upon rapamycin treatment immediately triggers the detachment of SYT1-GFP from the PM and its diffusion within the bulk ER network. Within this time frame of experimental manipulation, it is unlikely that PI4P depletion strongly impacts the pool of PI(4,5)P_2_ at the PM. Furthermore, PI(4,5)P_2_ depletion has no impact on SYT1-GFP localization. Given that PI4P dephosphorylation perturbed the PM association of both SYT1 and NET3C, our results suggest that PI4P acts a master lipid regulator for the localization of key ER-PM contact site proteins in *Nicotiana benthamiana* leaves.

**Fig. 1:**
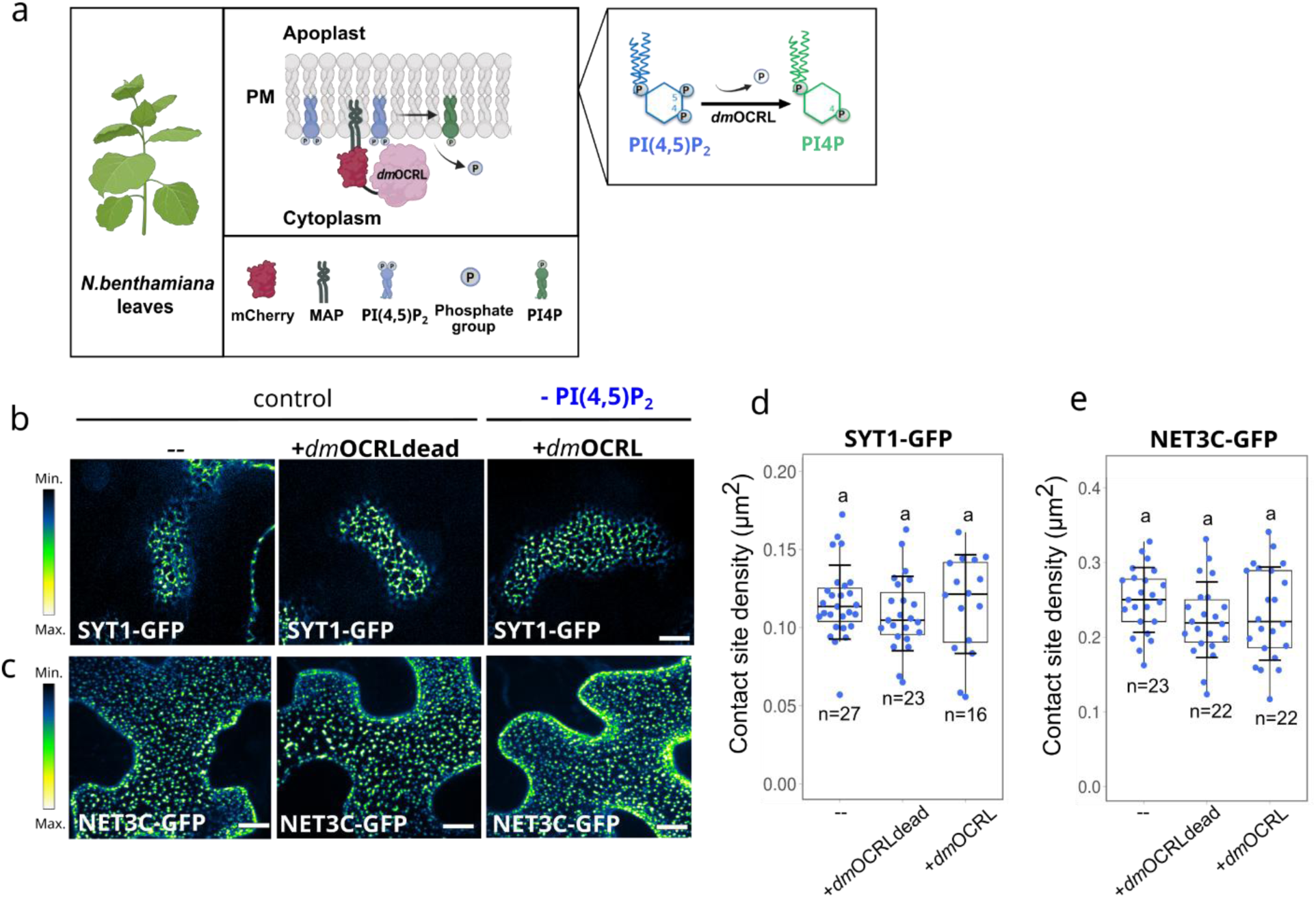
PI(4,5)P2 is dispensable for the establishment of ER-PM contact sites in *Nicotiana bentamiana* leaves. **(a)** Schematic representation of the iDePP system for plasma membrane PI(4,5)P_2_ depletion by *dm*OCRL. Created with BioRender.com. **b-c** Cortical view of leaf epidermal cell expressing **(b)** SYT1-GFP and maximum intensity projection of leaf epidermal cell expressing **(c)** NET3C-GFP co-expressed with mCherry-*dm*OCRLdead (middle panels) or mCherry-*dm*OCRL (right panels), respectively. **d-e** Quantification of **(d)** SYT1-GFP and **(e)** NET3C-GFP contact-site density. Horizontal lines represent the medians, whereas error bars represent SD. Data were analyzed using a one-way ANOVA test. No significant differences were observed among groups (p > 0.05). Each dot represents an individual cell, and n refers to the total number of cells analyzed from at least 5 plants per group. Scale bars= 10 µm.

**Fig. 2:**
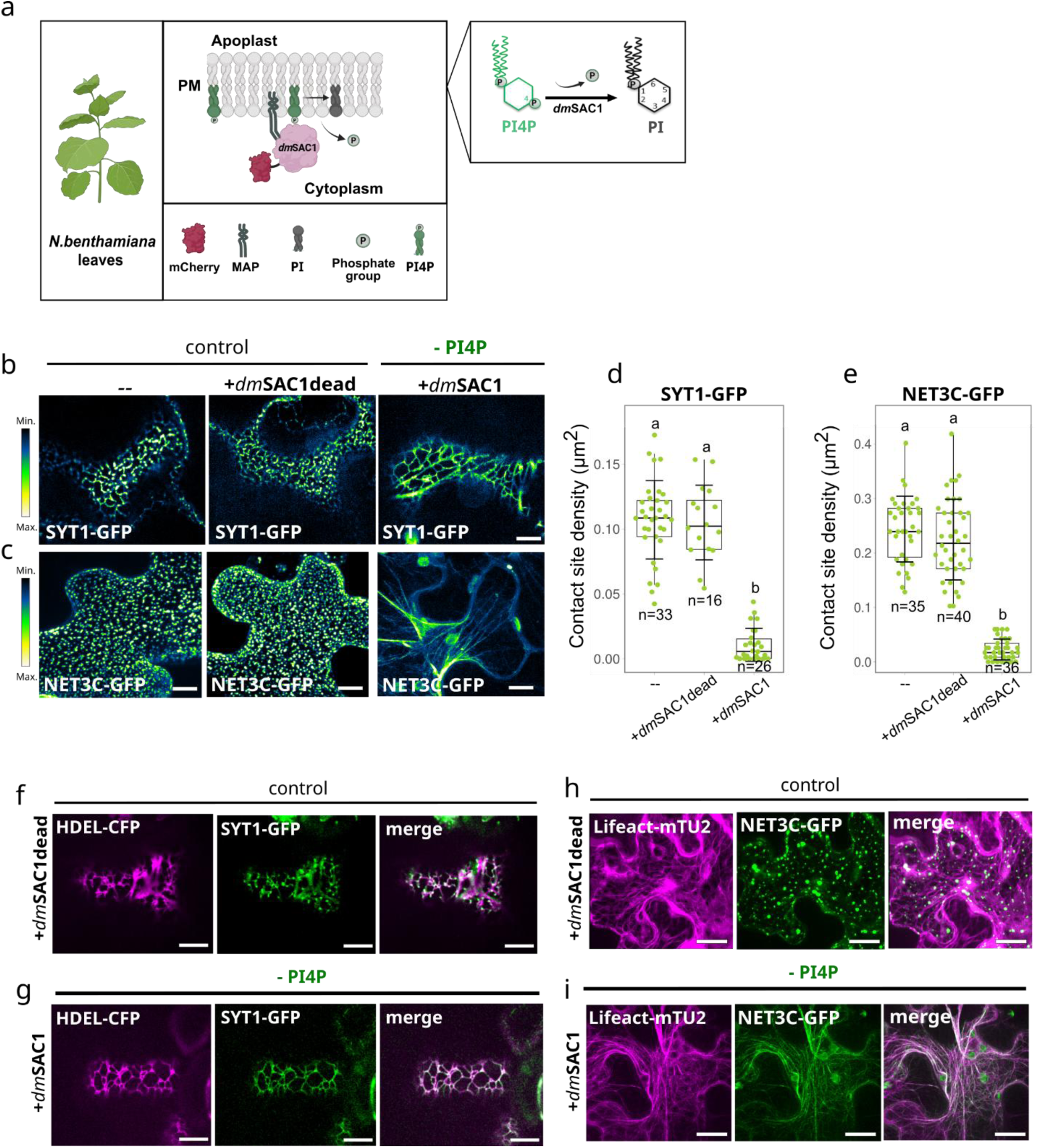
The plasma membrane localization of ER-PM contact site proteins is dependent on PI4P in *Nicotiana benthamiana* leaves. **(a)** Schematic representation of genetically-encoded system for plasma membrane PI4P depletion by *dm*SAC1. Created with BioRender.com. **b-c** Cortical view of leaf epidermal cells expressing **(b)** SYT1-GFP and maximum intensity projection of leaf epidermal cells expressing **(c)** NET3C-GFP co-expressed with *dm*SAC1dead-mCherry (middle panels) or *dm*SAC1-mCherry (right panels), respectively. **d-e** Quantification of **(d)** SYT1-GFP and **(e)** NET3C-GFP contact-site density. Horizontal lines represent the medians, whereas error bars represent SD. Letters denote statistically different groups calculated by Kruskal-Wallis test followed by Dunn’s post hoc test with adjusted p values (p < 0.01). Each dot represents an individual cell, and n refers to the total number of cells analyzed from at least 5 plants per group. **f-g** Cortical view of leaf epidermal cells expressing SYT1-GFP, HDEL-CFP and **(f)** *dm*SAC1dead-mCherry or **(g)** *dm*SAC1-mCherry. **h-i** Maximum intensity projection of leaf epidermal cell expressing NET3C-GFP, Lifeact-mTurquoise2 and **(h)** *dm*SAC1dead-mCherry or **(i)** *dm*SAC1-mCherry. Scale bars= 10 µm.

**Fig. 3:**
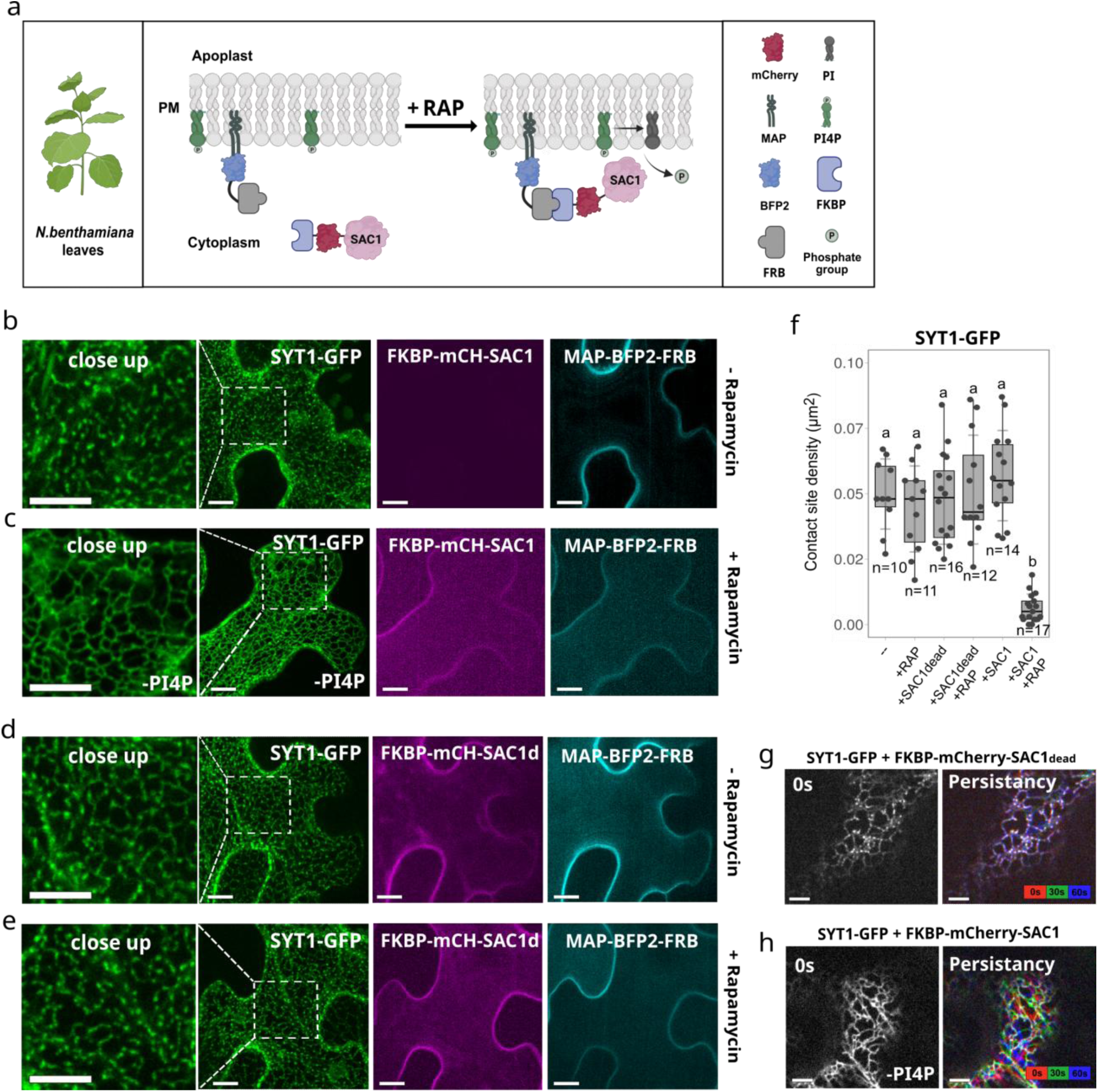
SYT1 contact sites detach rapidly from the plasma membrane in response to PI4P depletion. **(a)** Schematic representation of the rapamycin (RAP)-inducible system for plasma membrane PI4P depletion by yeast SAC1. Created with BioRender.com. **b-e** Maximum intensity projected images of epidermal *Nicotiana benthamiana* leaf cells transiently expressing the SYT1-GFP (left panels), the FKBP-mCherry-tagged **(b-c)** active or **(d-e)** catalytically dead yeast SAC1 (middle panels) and the PM anchor MAP-BFP2-FRB (right panels). In the absence of rapamycin, SYT1-GFP localizes in the contact sites and bulk ER (**b** and **d,** left panels). **(c)** Upon rapamycin treatment, SAC1 localizes at the PM and localization of SYT1-GFP is shifted from contact site to the bulk ER. **(e)** Rapamycin inducible PM targeting of SAC1dead did not impacted the localization of SYT1-GFP. **(f)** Statistical analysis of contact site density of SYT1-GFP represented in **(b-e)**. Horizontal lines represent the medians, whereas error bars represent SD. Each dot represents an individual cell, and n refers to the total number of cells analyzed from at least 5 plants per group. Letters show statistically different groups calculated by Kruskal-Wallis test followed by Dunn’s post hoc test with adjusted p values (p < 0.01). **g-h** The cell cortex of *Nicotiana benthamiana* leaf cells expressing the SYT1-GFP and **(g)** FKBP-mCherry-SAC1dead and **(h)** FKBP-mCherry-SAC1 upon rapamycin treatment. The images on the left represent the first time-lapse image (time point = 0s). The images on the right represent merged pseudo colored images of a 1 minute time-lapse. Scale bars =10 µm.

### PI4P is crucial for the formation of SYT1-containing contact sites in Arabidopsis root cells

Our transient expression experiments in *Nicotiana benthamiana* revealed that PI4P is essential for the localization of ER-PM contact site proteins, whereas PI(4,5)P_2_ is dispensable to their PM association. Next, we thought to validate these observations in a second plant species, different cellular system and stably transformed organism. To this end, we turned to *Arabidopsis thaliana* and focused on the regulation of SYT1 localization. As a first approach, we targeted PI4Kα, the enzyme that phosphorylates phosphatidylinositol into PI4P at the PM in Arabidopsis^43^. However, we previously showed that PI4Kα is essential in plants as *pi4k*α knockout mutants are gametophytic lethal. To bypass this lethality, we designed a tissue-specific CRISPR strategy. To this end, we used multiple guides RNA targeting the *PI4Kα* gene and expressed *CAS9* under the control of the *PIN-FORMED2* (PIN2) promoter that is active in epidermis and cortex in root cells, early during root development^44^ (**Fig.4a**). These *PIN2p>>PI4Kα* CRISPR mutant plants (hereafter *pi4kα crispr*) developed significantly shorter roots than the control plants and had impaired root hair development, in accordance with the key function of PI4Kα in root development^45^ (**Supplementary Figs.3a-d**). Next, we transformed complemented *syt1* mutants expressing SYT1p:SYT1:GFP with our *pi4kα crispr* construct and followed the localization of SYT1-GFP in root meristematic epidermal cells where the *pi4kα crispr* cassette was active. In control conditions, SYT1-GFP localized mainly in the cortical regions of root meristem cells, in accordance with its ER-PM tethering function, and to a lower extent in the perinuclear region (**Fig.4b, left panel**). Consequently, SYT1-GFP presents a high cortical index (i.e. ratio of fluorescence intensity between the apical/side cell region and the perinuclear region — see Materials and Methods) **(Fig.4c)**. Such high cortical index is typical of ER-PM contact site proteins and differs from proteins that localize to the bulk ER, such as the luminal ER marker HDEL-GFP, that accumulates more prominently in the perinuclear region of the ER network^31^ (**Fig.4b. middle panel**). In contrast to the control condition, SYT1-GFP accumulated in the perinuclear region of *pi4kα crispr* epidermal cells (**Fig.4b, right panel**). Consequently, SYT1-GFP exhibited a low cortical index in *pi4kα* mutant cells that was similar to the bulk ER marker HDEL-GFP (**Fig.4c**). Moreover, time-lapse imaging of the cortical region in differentiated root cells revealed that SYT1-GFP contact sites remain stable over a 50-second time lapse in control condition, as visualized by kymograph analyses and persistency mapping (**Fig.4d**). Conversely, in *pi4kα crispr* mutant, most of the SYT1-GFP localized in the dynamic ER (**Fig.4e**). Altogether, these results indicate that upon *pi4kα* mutation in the root epidermis and cortex, SYT1-GFP contact site localization is drastically affected, further supporting the importance of PI4P in this process.

Next, we investigated SYT1-GFP localization after inducible PI4P depletion in root cells using an orthogonal system different from the CRISPR strategy. To this end, we generated dexamethazone (dex) inducible MAP-*dm*SAC1-mCherry and MAP-*dm*SAC1dead-mCherry stable transgenic lines (**Fig.4f**). First, we tested the ability of the system to efficiently deplete PI4P by crossing these lines with the PI4P biosensor mCitrine-P4M^SidM^ and followed its localization in root meristematic cells. Upon 16h of dex induction, PI4P biosensor re-localized from the PM to the TGN/endosomes only when co-expressed with MAP-*dm*SAC1, but not MAP-*dm*SAC1dead (**Supplementary Fig.2d**). Next, we crossed the SYT1-GFP line with MAP-*dm*SAC1-mCherry and MAP-*dm*SAC1dead-mCherry and investigated the localization of SYT1-GFP in root meristematic cells. In the absence of dex, MAP-*dm*SAC1-mCherry and MAP-*dm*SAC1dead-mCherry were not expressed (**Figs.4g-h, upper left panels**) and SYT1-GFP localized in the cortical region of root meristem cells (**Figs.4g-h, upper right panels**). Upon 16h of dex induction, both enzymes were efficiently targeted to the membrane (**Figs.4g-h, down left panels**). SYT1-GFP localization was similar as in the control condition when coexpressed with the inactive SAC1 (**Fig.4g down right panel**). However, when co-expressed with the active SAC1, SYT1-GFP localization shifted towards the perinuclear region of the root meristematic cell (**Fig.4h down right panel**). Quantification analyses showed that SYT1-GFP cortical index was significantly reduced only when SYT1-GFP was co-expressed with the MAP-*dm*SAC1-mCherry (**Fig.4i**). Again, this analysis is consistent with a key function of PI4P for the localization of SYT1-GFP at ER-PM contact sites.

It was previously suggested that SYT1 localization in contact sites might depend on the presence of PI(4,5)P_2_ in the PM in Arabidopsis^17^, based on the correlation between the amount of SYT1-GFP-containing contact sites and the accumulation of a PI(4,5)P_2_ biosensor. However, it was never formally demonstrated whether PI(4,5)P_2_ is indeed required for this localization. To test the impact of PI(4,5)P_2_ depletion on SYT1-GFP localization, we employed our dex inducible iDePP system that selectively depletes PI(4,5)P_2_ in many cell types in Arabidopsis, including the root^35^ (**Supplementary Fig. 4a**). We crossed the SYT1-GFP reporter line with dex inducible MAP-mCherry-OCRL line and MAP-3xmCherry line, that we used as control. We then followed the localization of SYT1-GFP in meristematic root cells 16h after dex treatment, a treatment duration that allowed full solubilization of PI(4,5)P_2_ biosensor in root cells^35^. In the absence of dex, MAP-3xmCherry and MAP-mCherry-OCRL were not expressed (**Supplementary Figs. 4b-c, upper left panels**) and most of the SYT1-GFP localized in the cortical region of root meristem cell (**Supplementary Figs. 4b-c, upper right panels**). Upon dex treatment, both MAP-mCherry-OCRL and MAP-3xmCherry were expressed and localized to the PM (**Supplementary Figs. 4b-c, down left panels**). However, SYT1-GFP remained localized in the cortical region of the cell, even when co-expressed with the active enzyme (**Supplementary Figs. 4b-c, down right panels**). Statistical analysis of SYT1-GFP cortical index did not show any difference among different groups (**Supplementary Fig. 4d**).

Altogether, these results indicate that under basal conditions, PI4P is crucial for the localization of SYT1-GFP in ER-PM contact sites in both Arabidopsis root cells and *Nicotiana benthamiana* leaves, while PI(4,5)P_2_ does not play an important role in this process.

### The PI4P phosphatase SAC7 associates with SYT1-containing contact sites in root hair cells

While the above analysis demonstrated that PI4P is required for SYT1 localization at ER-PM contact sites, we next wondered whether the local regulation of PI4P levels at the PM might regulate SYT1 localization and its underlying function. As a first approach, we investigated whether regulators of PI4P metabolism might be present within the SYT1 interactome (see companion manuscript – Morello-López et al. 2026). Among the SYT1 interactors, we found SAC7 (also known as RHD4), a PI4P phosphatase^29^, as highly enriched in AP-MS experiments (**Fig.5a**). SAC7 was classified as a dual-localized protein at the ER and PM in LOPIT analysis, and it was also detected in Turbo-ID proximity labeling experiments, but with a lower enrichment score (**Fig.5a**). Our previous work demonstrated that the PI4P phosphatase SAC7 regulates PI4P levels at the PM within plasmodesmata, thereby controlling MCTP4 docking and cell-to-cell diffusion^31^. The mechanism of SYT1 targeting to the PM is analogous to that of MCTP4 — they both contain C2 lipid binding domains through which they bind anionic lipids^16,46^. Altogether, SYT1 proteomic data as well as the known role of SAC7 at plasmodesmata contact sites suggests that SAC7 might function within SYT1 contact sites. First, we verified that SYT1 and SAC7 might form a complex *in planta*. To test this hypothesis, we performed co-immunoprecipitation in Arabidopsis seedlings. We used anti-mCherry antibodies to immunoprecipitate 2xmCherry-SAC7, or the PM marker Lti6b-2xmCherry as a control, and detected the presence of SYT1 using specific anti-SYT1 antibodies. We found that 2xmCherry-SAC7 but not Lti6b-2xmCherry co-immunoprecipiated SYT1, confirming that the two proteins can be found in the same complex *in planta* (**Fig.5b**).

This prompted us to investigate whether SAC7 might be associated with SYT1-containing contact sites. To test this, we crossed SYT1p:SYT1:GFP/*syt1* and SAC7p:2xmCherry:SAC7/*sac7* transgenic lines (hereafter SYT1-GFP and mCherry-SAC7, respectively) and followed their subcellular localization. Previous studies, including our own, have shown that SAC7 is essential for root hair growth and accumulates in trichoblast cells and growing root hairs^29–32^.

We found that similarly to mCherry-SAC7, SYT1-GFP also accumulated in the root hair bulge and at the tip of the growing root hairs (**Supplementary Fig. 5a**). We thus investigated the subcellular localization of mCherry-SAC7 and SYT1-GFP in detail in these cells. Using Airyscan super-resolution confocal microscopy, we found that mCherry-SAC7 localized in punctate structures in the cortical region of trichoblast cells that were often closely associated with SYT1-GFP-containing ER-PM contacts and even sometimes overlapped (**Figs.5c-d**). Moreover, the punctate localization of SAC7 was especially pronounced in the root hair bulge, which was not the case for the ER bulk marker HDEL-RFP (**Supplementary Fig. 5b**). The root hair bulge is the area where root hairs initiate in trichoblasts but are not actively growing yet. We used two-color total internal refection fluorescence microscopy (TIRF) microscopy to follow colocalization and dynamics of SYT1-GFP and mCherry-SAC7 at the PM in the root hair bulge. While SYT1-GFP exhibited stable localization within established contacts, the localization of mCherry-SAC7 was notably dynamic: mCherry-SAC7 transiently associated with SYT1-GFP before dissociating, a process that could recur over time (**Figs.5e-f**). Furthermore, the recruitment of mCherry-SAC7 to SYT1-GFP contact sites was sometimes associated with the fading away or even disappearance of SYT1-GFP signal (**Fig.5f, black arrows**). However, in many cases, the recruitment of mCherry-SAC7 to SYT1-GFP-containing contact sites was transient and the SYT1-GFP signal increased again after the departure from mCherry-SAC7 (**Fig.5f, red arrows**). This data suggested that recruitment of SAC7 to SYT1-containing ER-PM contacts might transiently displace SYT1 from the PM through local depletion of PI4P. This might explain why at a given time point most mCherry-SAC7 is associated with SYT1-GFP but does not directly colocalize. We thus decided to analyze their colocalization over time.

To this end, we quantified the colocalization between mCherry-SAC7 and SYT1-GFP using the DiAna ImageJ tool (see Materials and Methods). This analysis revealed that around 60% of SYT1-GFP puncta associated with mCherry-SAC7 and vice versa within a 40-seconds time frame (**Fig.5g and Supplementary Figs.5c-d**).

Collectively, these data demonstrate that SAC7 and SYT1 are present in the same complex and that SAC7 exhibits a dynamic association with SYT1-containing contact sites in trichoblasts, notably in non-growing bulging root hairs.

### SYT1 contributes to root hair growth

It has been shown that SAC7 is highly expressed in trichoblasts in Arabidopsis, and plants lacking SAC7 develop short root hairs in comparison to the control^29–31^. Because the most salient phenotype of the *sac7* mutant is short root hairs, we next investigated the subcellular localization of SYT1-GFP and mCherry-SAC7 in actively growing root hairs. When root hairs were growing, the localization of SYT1-GFP and mCherry-SAC7 was different than in bulging root hairs. Indeed, we found that both proteins accumulated and colocalized inside the root hair close to the growing tip (**Fig.6a**). This localization pattern is reminiscent of the ER membrane marker, which is also present at the tip of root hairs (**Supplementary Fig.6**). We also noticed that SYT1 contact sites were mostly found in the shank region of the growing root hair (**Figs.6a-b, left panels**). Unlike in bulging root hairs, mCherry-SAC7 displayed ER-like localization in growing root hairs and only rarely associated with SYT1-GFP contact sites in the shank region (**Figs.6a-b**). Next, we tested if SYT1 and SAC7 are genetically linked. We crossed *sac7-3 with syt1-2* (hereafter *sac7* and *syt1*, respectively) mutant plants and focused on the root hair phenotype of single and double mutants. We also included *syt1syt3* double mutants in our analysis as *SYT3* is functionally redundant to *SYT1*^6,47^. Our results showed that unlike *sac7* mutant plants, *syt1* single mutants didn’t exhibit root hair phenotype. However, when both SYT1 and SYT3 were mutated, the plants developed significantly shorter root hairs in comparison to the control, although not as strong as *sac7* plants (**Fig.6c**). The *sac7syt1* double mutant and *sac7syt1syt3* triple mutants exhibited an additive decrease in root hair length relative to the *sac7* single mutant, suggesting that SAC7 and SYT1 contribute to root hair elongation through partially independent or parallel pathways. Interestingly, the *sac7syt1* double mutant and *sac7syt1syt3* triple mutant showed a significant reduction in root hair density compared to either single mutant, indicating that both SAC7 and SYT1 are required for proper root hair establishment (**Figs.6d-e**). Together, these results suggest that SAC7 and SYT1 cooperatively regulate root hair formation and elongation, with SAC7 playing a predominant role.

**Fig. 4:**
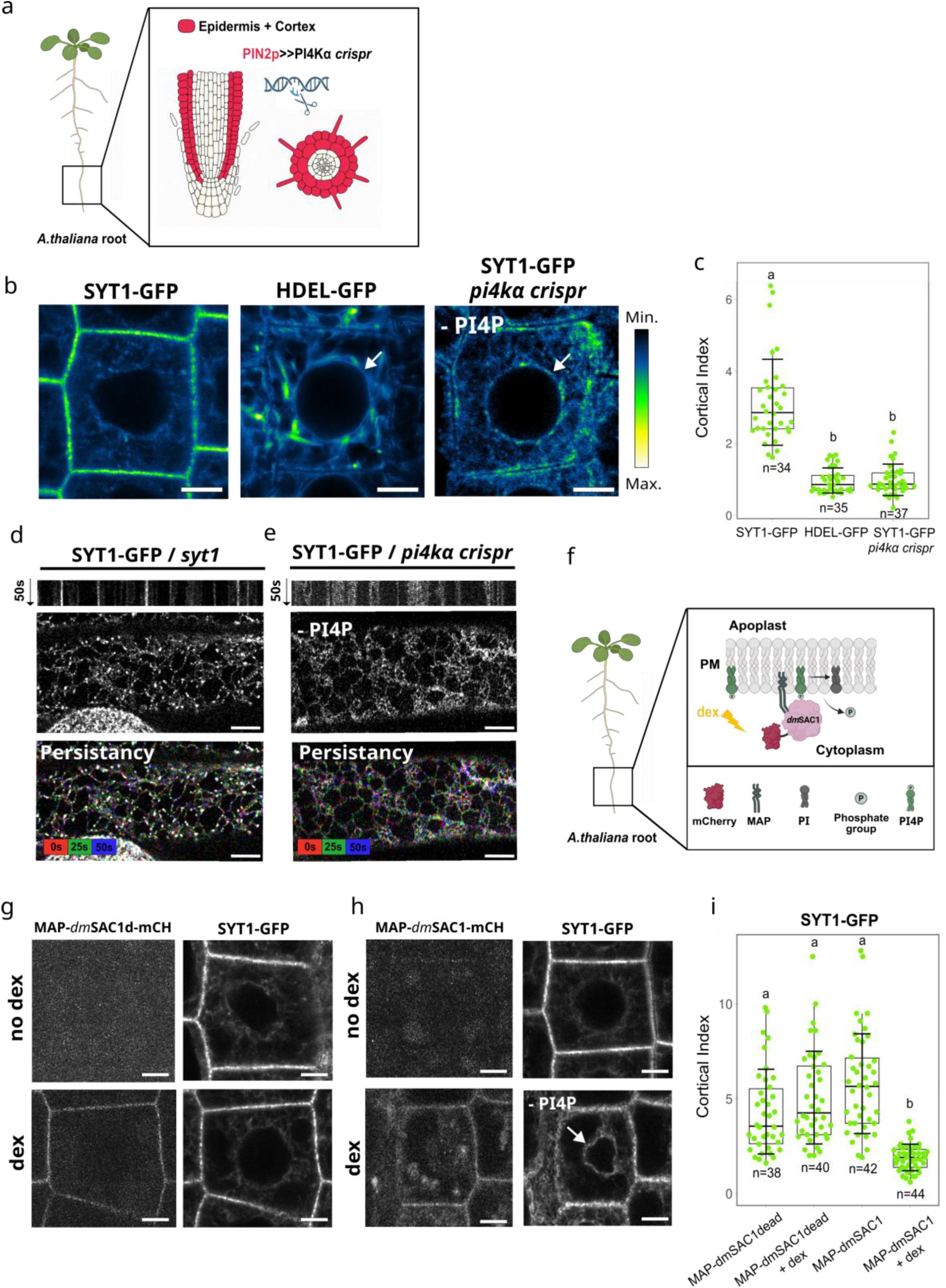
The plasma membrane localization of SYT1 is dependent on PI4P in Arabidopsis thaliana roots. **(a)** Schematic representation of the tissue-specific CRISPR strategy for PI4P depletion in Arabidopsis roots. **(b)** Representative images of root meristematic epidermal cells expressing SYT1p:SYT1:GFP in *syt1*, HDEL-GFP and SYT1p:SYT1:GFP in *pi4kα crispr* mutant. The arrows indicate the localization of HDEL-GFP and SYT1p:SYT1:GFP in the perinuclear region. Scale bars = 5 µm. **(c)** Statistical analysis of cortical index of **(b)**. Horizontal lines represent the medians, whereas error bars represent SD. Each dot represents an individual cell, and n refers to the total number of cells analyzed from at least 5 seedlings per group. Different letters indicate statistically significant differences between groups calculated by Kruskal-Wallis test followed by Dunn’s post hoc test with adjusted p values (p < 0.01). **(d-e)** Cortical view of differentiated root epidermal cells expressing SYT1p:SYT1:GFP in **(d)** *syt1* and in **(e)** *pi4kα crispr* mutant with their respective kymographs. The images in down panels represent merged pseudo colored images of a 50s time-lapse. Scale bars =5 µm. **(f)** Schematic representation of the dexamethasone (dex) inducible plasma membrane PI4P depletion by *dm*SAC1 in Arabidopsis cells. Created with BioRender.com. **(g)** Representative images of the fluorescent signal corresponding to MAP-*dm*SAC1dead-mCherry (left upper panel) and SYT1p:SYT1:GFP (right upper panel) in the same cell, without dex treatment, and MAP-*dm*SAC1dead-mCherry (left down panel) and SYT1p:SYT1:GFP (right down panel) in the same cell after 16 hours of dex treatment. **(h)** Representative images of the fluorescent signal corresponding to MAP-*dm*SAC1-mCherry (left upper panel) and SYT1p:SYT1:GFP (right upper panel) in the same cell, without dex treatment, and MAP-*dm*SAC1-mCherry (left down panel) and SYT1p:SYT1:GFP (right down panel) in the same cell after 16 hours of dex treatment. The arrow indicates the localization of SYT1p:SYT1:GFP in the perinuclear region. **(i)** Statistical analysis of SYT1p:SYT1:GFP cortical index of **(g)** and **(h).** Scale bars = 5 µm.

**Fig. 5:**
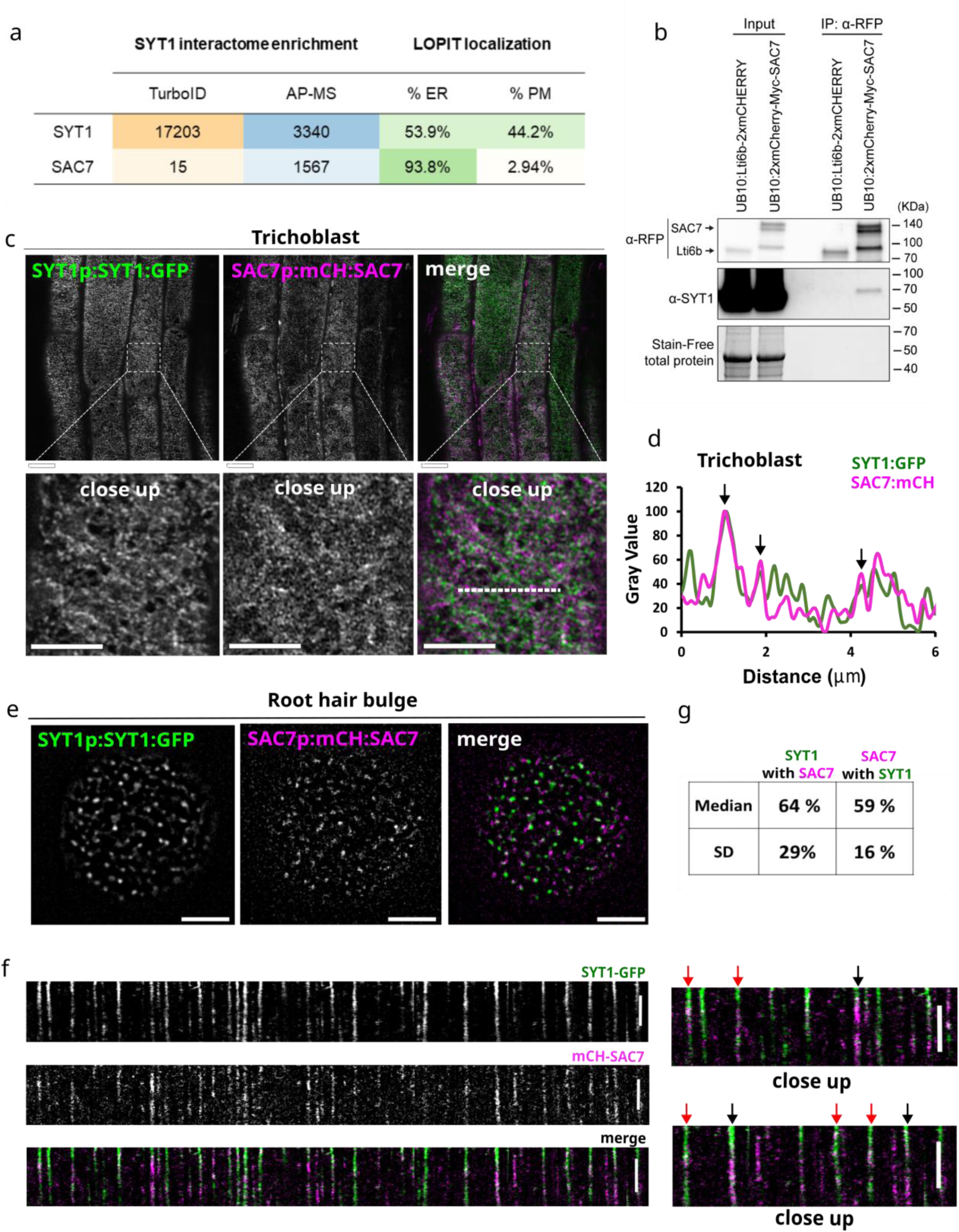
SAC7 transiently associates with SYT1 in non-growing root hair bulges. **(a)** The Affinity Enrichment Scores (AES) of SYT1 and SAC7 peptides from Turbo-ID and AP-MS proteomic experiments and percentage of SAC7 and SYT1 protein distributions at the ER and PM using Localization of Organelle Proteins by Isotope Tagging (LOPIT) proteomic approach (see companion manuscript – Morello-López et al., 2026). **(b)** Co-immunoprecipitation of SAC7 with SYT1. Arabidopsis seedlings expressing Lti6b-2xmCherry, and 2xmCherry-SAC7 were pulled down using anti-RFP magnetic beads. The interaction with SYT1 was tested using anti-SYT1 antibodies**. (c)** Cortical view of a trichoblast cell expressing SYT1-GFP and mCherry-SAC7. Scale bars = 5 µm. **(d)** Fluorescence intensity plot of the signals in **(c)** along the white dashed line. The arrows indicate overlapping fluorescence intensities of SYT1-GFP and mCherry-SAC7. **(e)** Representative images of a root hair bulge expressing SYT1-GFP and mCherry-SAC7 taken by the TIRF microscope. Scale bars = 5µm. **(f)** Kymographs from 40-second time-lapses of root hair bulges, showing individual SYT1-GFP and mCherry-SAC7 puncta. Black arrows indicate disappearance of SYT1-GFP signal upon mCherry-SAC7 recruitment to the contact site. Red arrows indicate transient recruitment of mCherry-SAC7 to the SYT1-GFP contact sites. Scale bar = 20s. **(g)** Quantification analysis of colocalization between SYT1-GFP and mCherry-SAC7 puncta of **(e)**.

**Fig. 6:**
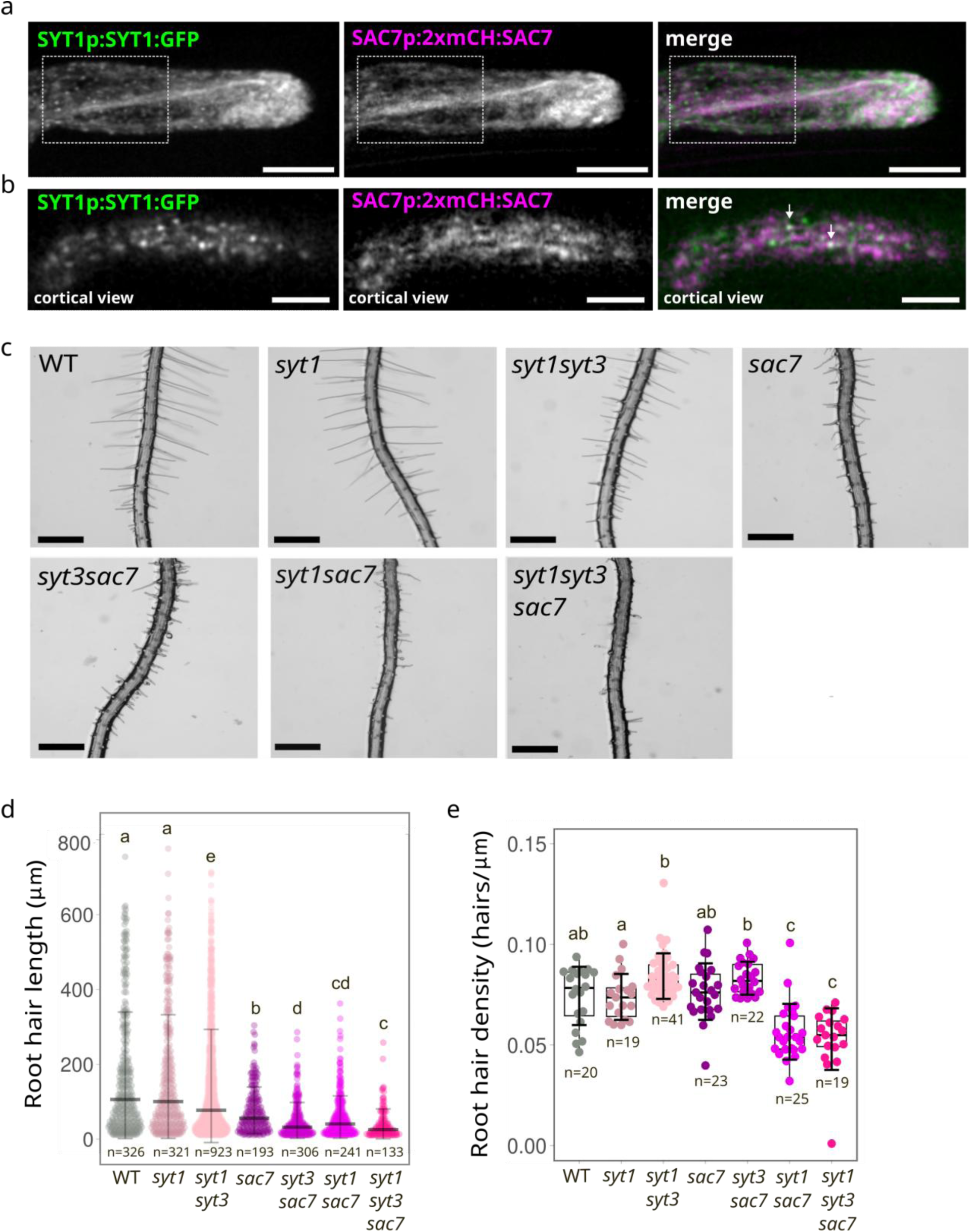
SYT1 is involved in root hair growth. **(a)** Maximum intensity projection of a root hair expressing SYT1-GFP and mCherry-SAC7. Dashed boxes indicate the shank region of a root hair. Scale bars = 10 µm **(b)** Cortical view of shank regions of root hairs presented in **(a)**. The arrows indicate SYT1-GFP and mCherry-SAC7 association in the contact sites. Scale bars = 5 µm. **(c)** Representative images of Arabidopsis root hair phenotypes in different mutant backgrounds. **d-e** Quantification of the root hair length **(d)** and root hair density **(e)** of the images shown in **(c)**. Horizontal lines represent the medians, whereas error bars represent SD. Each dot represents an individual root hair **(d)** or individual root **(e)**. Differences were assessed using a Kruskal–Wallis test followed by Dunn’s post hoc test with adjusted p values (p < 0.05). Groups with different letters are significantly different. Scale bars = 500 µm.

### SAC7 inhibits the establishment of SYT1-containing contact sites to allow root hair growth

Having established that both SYT1 and SAC7 are important for root hair growth, we sought to understand the mechanism driving this process. It was previously published that *sac7* mutant plants have elevated levels of PI4P in comparison to the control^29,30^. Since we showed that PI4P is important for SYT1 tethering to the PM, we hypothesized that SAC7 might regulate SYT1 localization through regulation of PI4P levels at the PM. To test this, we crossed SYT1p:SYT1:GFP with *sac7* mutant plants (SYT1-GFP/*sac7*). As a control, we used a wild-type sister plant from the same cross (SYT1-GFP/wt). In control, most of the SYT1-GFP accumulated in the bulk ER close to the tip of growing root hairs, while the stable contacts were visible away from the tip, mostly in the shank region of root hairs (**Fig.7a and Supplementary video.1**). In contrast, in *sac7* mutant, SYT1-GFP contact sites ectopically accumulated at the tip of growing root hairs (**Fig.7b and Supplementary video.2**). Similarly, we found that SYT1-GFP accumulated at the tip in mature root hairs of wild-type plants, at a stage where they stopped growing (**Supplementary Fig.7 and Supplementary video.3**).

**Fig. 7.**
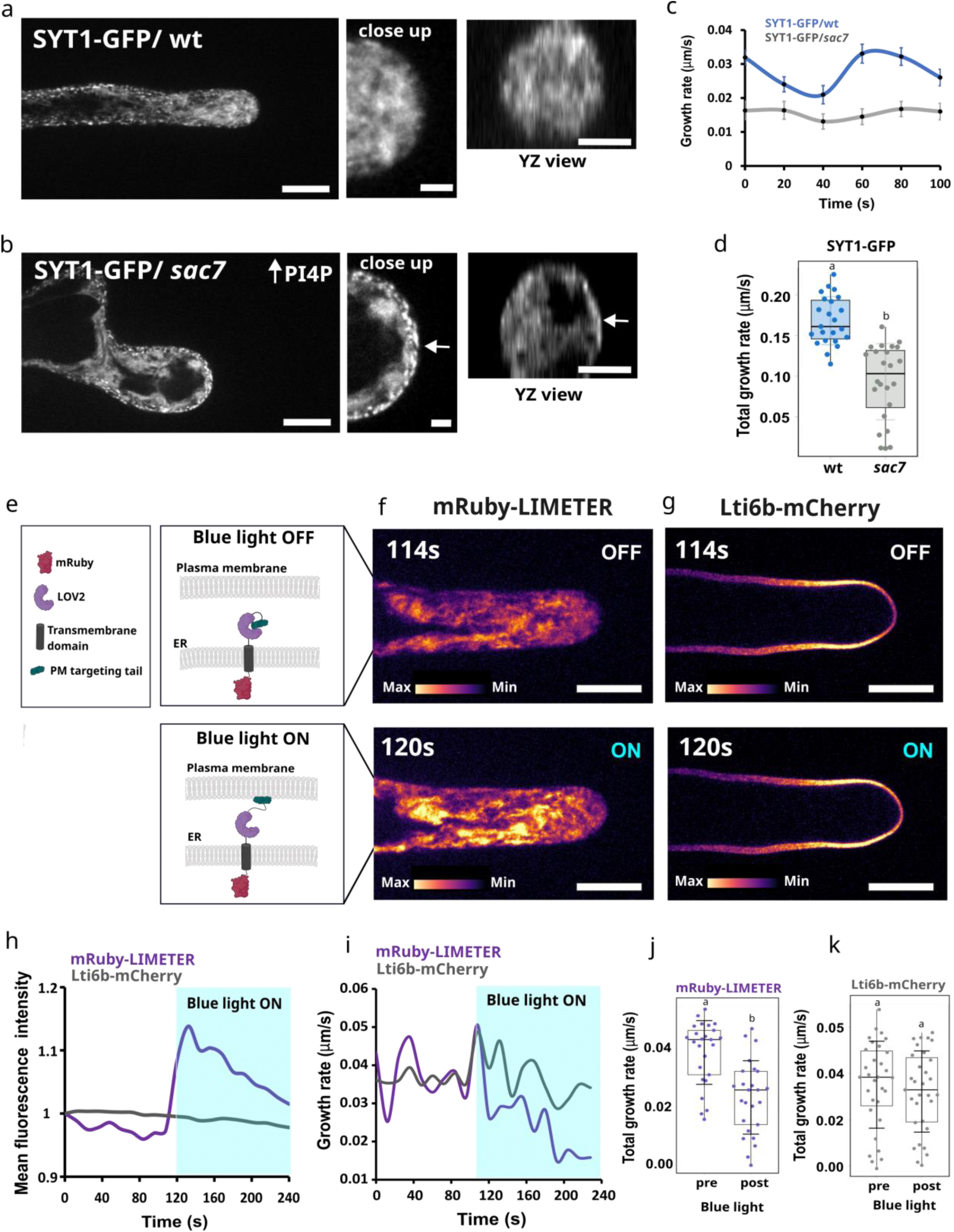
SAC7 inhibits formation of SYT1 contact sites at the root hair tip. a-b. Maximum intensity projection of a root hair expressing SYT1-GFP in **(a)** wt and **(b)** *sac7* background. Scale bars=10 µm. The arrows on the close up and YZ view images indicate accumulation of SYT1-GFP at the root hair tip. Scale bars=5 µm. **c-d** Quantification of **(c)** root hair growth rate per time point during 2-minute time lapse and 20-second interval. Dots represent mean growth rates whereas error bars represent SE per each time point. **(d)** Statistic analysis of maximal growth rate of SYT1-GFP in wt and *sac7* root hairs during 2-minute time lapse. Horizontal lines represent the medians whereas error bars represent SD. Each dot represents an individual root hair and n refers to the total number of analyzed root hairs. Different letters indicate statistically significant differences between groups (p < 0.01). **(e)** The schematic representation of the LiMETER optogenetic system. Created with BioRender.com. **f-g** Representative images of root hairs expressing **(f)** mRuby-LiMETER and **(g)** Lti6b-mCherry in resting state (upper panels) and after blue light activation (down panels). Scale bars=10 µm. **(h)** Plot of mean fluorescence intensities, normalized to 1, from root hairs expressing mRuby-LIMETER and Lti6b-mCherry, acquired by time-lapse imaging at 12-second intervals over a 4-minute period. **(i)** Growth rate plots from 4-minute time-lapse imaging of root hairs expressing mRuby-LiMETER and mCherry-Lti6b. Frames were captured at 12-second intervals. Blue light was applied at 2-minute time point. **j-k** Quantification of the maximal growth rate of root hairs expressing **(j)** mRuby-LiMETER and **(k)** Lti6b-mCherry pre and post blue light application. Horizontal lines represent the medians, whereas error bars represent SD. Each dot represents an individual root hair, and n refers to the total number of analyzed root hairs. Letters show statistically different groups calculated by Kruskal–Wallis test followed by Dunn’s post hoc test (p< 0.01).

The short root hairs in the *sac7* mutant could result from either normal growth that stops prematurely—such as through early activation of a cell death program^48^—or from a slower growth rate compared to the wild type. To distinguish between these possibilities, we took 2-minute time lapses of SYT1-GFP/wt and SYT1-GFP/*sac7* root hairs and measured their respective growth rates. We observed that in 2-minute time frame, *sac7* mutants retained SYT1-stable contact sites at the root hair tip, correlating with a significant reduction in root hair growth rate in comparison to control (**Figs.7c-d**). Together, these data indicate that loss of SAC7 enhances SYT1 tethering to the PM, which might in turn limit the growth rate of root hairs.

SYT1 forms stable ER-PM contacts in bulging root hairs, which pause their growth as they prepare for root hair outgrowth. By contrast, SYT1-containing contact sites are excluded from the tip of growing root hair in the wild type, but populate the tip in slow growing *sac7* or mature wild-type root hair. Together, these data suggest that the establishment of stable ER-PM contacts might negatively impact polarized growth.

However, so far, we found a correlation between the presence of SYT1-containing ER-PM contacts at the tip and the restriction of growth, but not a causal relationship. To challenge this hypothesis and demonstrate that slow root hair growth can be triggered by ectopic ER-PM contact sites at the tip, we turned to an optogenetic strategy aimed at rapidly increasing ER-PM tethering by using blue light. To this end, we employed the LiMETER (Light-inducible Membrane-Tethered cortical ER)^49^ optogenetic system to acutely induce ER-PM tethering in growing root hairs. LiMETER comprises an ER transmembrane domain, an mRuby fluorescent tag in the ER lumen, a light-switchable LOV2 domain derived from *Avena sativa* phototropin 1, and a PM-targeting motif consisting of a polybasic tail that binds to negatively charged phospholipids at the PM. In its resting state, the PM-targeting motif is caged within the LOV2 domain, but upon blue light activation, the motif is released and binds to the PM, thereby inducing ER-PM tethering (**Fig.7e**). It was previously shown that LiMETER can specifically and reversibly induce ER-PM tethering in Arabidopsis upon blue light activation in different cell types^49^. To test if the system is working in our conditions, we first aimed to induce LiMETER PM tethering in Arabidopsis root differentiated cells. In resting state, LiMETER localized in the bulk ER network in the cortical region of differentiated root epidermal cells. Upon blue light activation, LiMETER was recruited to the ER-PM contacts within seconds (**Supplementary video.4**). Next, to investigate the effect of induced ER-PM tethering in root hairs, we captured 4-minute time-lapse images of root hairs expressing LiMETER and applied blue light after 2 minutes. To account for potential phototoxicity, mCherry-Lti6b served as a negative control. LiMETER localized in rapidly remodeling ER when it was not activated by blue light (**Fig.7f, upper panel**), and showed a strong and rapid increase in fluorescence intensity seconds after blue light activation, unlike the Lti6b-mCherry control (**Figs.7f-g, and Supplementary videos.5-6**). We observed that blue light treatment triggered an immediate decrease in root hair growth, which was significant in comparison to the resting state (**Figs.7i-j**). Importantly, this effect was specific to LiMETER as Lti6b-mCherry fluorescence intensity and the growth rate of Lti6b-mCherry root hairs were not significantly different from the control condition without blue light (**Figs.7h-k**). Therefore, even a transient induction of ER-PM contact sites in root hairs was sufficient to significantly affects their growth rate. Altogether, our results suggest a causal role for ER-PM tethering in controlling polarized cell growth, whereby a high local density of contact sites restrict growth (i.e. in bulging root hair, *sac7* mutants or blue-light activated LiMETER), while SAC7-mediated depletion of contacts allows localized polar growth.

## Discussion

### PI4P is crucial for plasma membrane tethering of ER-PM contact site proteins in plants

Using different genetic tools for *in vivo* depletion of phosphoinositides, we showed that PI4P plays a key role in regulating the formation and dynamics of ER-PM contact sites in plants. These results show the importance of verifying *in vivo* protein-lipid interactions that have been studied *in vitro*. Indeed, early studies on Arabidopsis SYT1 clearly showed that these proteins interact with several phosphoinositides *in vitro*^15,16,25^.

However, because of the similarities between SYT1 and human E-SYTs, the field focused on PI(4,5)P_2_. This is surprising since PI4P is the major phosphoinositide at the plant PM, not PI(4,5)P_2_, and this lipid also interacts with SYT1^15,26,28,50^. Several studies described at great length the interaction between PI(4,5)P_2_ and SYT1 *in vitro*, showing that this interaction can induce membrane binding and even tethering and deformation. However, none of these studies included PI4P in their analysis and detailed lipid-binding properties of other plant ER-PM contact sites proteins remain unexplored.

Here, we show that PI4P rather than PI(4,5)P_2_ is a key lipid for SYT1 and NET3C plasma membrane localization *in vivo*. We previously showed that the electrostatic field of the plant PM is powered by PI4P rather than PI(4,5)P_2_, which likely explains the prominent role of PI4P in SYT1 and NET3C interaction with the PM^26^. Previous studies showed that SYT1 PI(4,5)P_2_ binding is mediated by a polybasic site in its C2A domain^25^. Such a site is likely to also bind to PI4P *in vitro* and also *in vivo*. In accordance, we recently showed that another class of C2 domain–containing proteins localized at plasmodesmata, multiple C2 domains and transmembrane region proteins (MCTPs), rely on PM PI4P enrichment for membrane tethering^31^.

On another hand, lipid binding properties of NET3C are less understood. However, NET3C contains several basic motifs (lysine/arginine-rich residues) at its C-terminus that may mediate its association with negatively charged PI4P. Consistent with this hypothesis, a NET3C mutant in which a C-terminal lysine residue was substituted with alanine exhibited reduced PM tethering and diminished localization to actin filaments^22^. In addition to SYT1 and NET3C, several newly identified ER–PM contact site proteins require PI4P for their PM localization (see companion manuscript – Morello-López et al., 2026). Therefore, PI4P-mediated PM tethering may serve as a common mechanism for the recruitment and anchoring of ER-PM proteins at the PM at least under basal conditions.

However, we cannot rule out that other anionic lipids, such as PI(4,5)P2 or PS, contribute to their localization in specific cell types or under stress conditions, as reported for other proteins that rely on anionic lipids for their PM localization^51–53^.

Indeed, it was previously shown that increasement of PI(4,5)P_2_ levels upon ionic stress caused cortical expansion of SYT1-containing contact sites in cotyledon epidermal cells^17^. Importantly, the *in vitro* studies performed on SYT1 C2 domains also detected a calcium-dependent interaction with PS^6,15,19,25^. Since calcium is often released upon stresses that also trigger ER-PM contact site rearrangements, it is possible that SYT1-PS interaction could be involved in the remodeling of SYT1 localization during stress episodes^54^.

In any case, PI(4,5)P_2_ is clearly dispensable for SYT1 and NET3C interaction with the PM under basal conditions, while PI4P is not.

### SAC7 regulates ER-PM contact site dynamics during root hair polar growth

Our results suggest that transient association of SAC7 with SYT1-containing contact sites negatively regulates SYT1–PM tethering. This feedback regulation operates during cell polarization of root hairs with opposite outcome linked to growth. In the bulge of root hairs, a specialized polar membrane domain in trichoblast where root hairs are preparing to grow but are not actively growing yet, SYT1-mediated ER-PM contacts are very prominent. There, SAC7 transiently associates with SYT1 leading to only a brief dissociation of SYT1 from the PM. By contrast, SAC7 massively accumulates at the tip of growing root hair, mostly in the bulk ER, constantly detaching SYT1 from the PM in this tip region. As a result, SYT1 is continuously tethered at the PM at the root hair tip in *sac7*, which correlates with their slow growth rates, suggesting that ER-PM contacts might act as breaks of polarized growth. Indeed, fast optogenetic upregulation of ER-PM contact sites almost immediately reduced root hair growth rate, suggesting a causal link between the absence of stable ER-PM contact sites and polarized growth.

The SAC7-dependent regulation of SYT1 contact sites is likely happening only at the tips of growing root hairs as stable SYT1-containing contact sites are found in the shank region of root hairs (**Fig.6a-b and Fig.7a**). Similarly, ER–PM contact sites in rod-shaped fission yeast accumulate at the non-growing lateral cortex and are excluded from actively growing cell tip. Moreover, overexpression of an artificial ER–PM tether in rod-shaped yeast cells, strongly affected cell polarity and secretion^55^. Likewise, stable ER–

PM contact sites in neurons are absent from regions of high secretory activity^56^. Therefore, we can speculate that the negative correlation between stable ER–PM contact site formation and polarized growth may represent an evolutionarily conserved feature.

In addition to stable ER-PM contact sites at the tip, other factors may contribute to the root hair phenotype observed in *sac7* mutant. For example, we cannot exclude the possibility that the phenotype results, at least in part, from altered vesicular trafficking to the root hair tip, as specific phosphoinositide signatures are known to be critical for polarized growth^52,57^. Furthermore, the altered PI4P levels in the *sac7* mutant may affect the localization and function of proteins other than ER–PM contact site components. Indeed, PI4P has recently been shown to regulate the assembly of ER-localized tail-anchored proteins, suggesting that additional PI4P-dependent processes could contribute to the *sac7* phenotype^32^.

An important question arising from our findings is the mechanism by which SAC7 regulates SYT1 contact-site dynamics during polar root hair growth. The ER localization of SAC7 suggests that PI4P consumption occurs at the ER and therefore likely depends on the transport of PI4P from the PM to the ER membrane. This model is consistent with the activity of SAC1, the animal ortholog of SAC7, which dephosphorylates PI4P in *cis*—that is, within the same membrane in which it resides—rather than in *trans*^58^.

Although the molecular machinery responsible for this lipid transfer remains largely uncharacterized in plants, members of the OSBP-RELATED PROTEIN (ORP) family are conserved and represent strong candidates for mediating PI4P trafficking at ER–PM contact sites^59,60^.

In this study we propose a negative-feedback mechanism in which SYT1-dependent tether formation facilitates the recruitment of SAC7, which in turn locally consumes PI4P and limits further tether stabilization. Such regulation may not be restricted to SYT1 alone. Indeed, we previously demonstrated a role for SAC7 in regulating cell-to-cell communication, where it controls MCTP4-mediated tethering within plasmodesmata in a cell type-specific manner^31^. Together, we propose that SAC7-mediated PI4P depletion could represent a general mechanism controlling the assembly and disassembly of diverse ER–PM contact sites in plants.

## Methods

### Plant material and growth conditions

*Nicotiana benthamiana* and *Arabidopsis thaliana* plants were used in this study. All *A.thaliana* plants are of *Columbia-0* (Col-0) ecotype. Mutant line *sac7-3* (SALK_079231) was obtained from J.Schiefelbein (Michigan University) and was characterized in a previous studies^29,31^. Mutant lines *syt1-2* (SAIL_775_A08), *syt3* (SALK_037933) and *syt1/syt3* double mutant were previously described^6,36^. Double and triple mutants *sac7/syt1* and *sac7/syt1/syt3* were obtained by crossing. The following stable transgenic lines were previously published and characterized: SYT1p:SYT1-GFP/*syt1*^36^, SAC7p:mCitrine:SAC7/*sac7* and SAC7p:2xmCherry:SAC7/sac7^31^, UBQ10p:GVG:MAP-mCherry:dmOCRL and UBQ10p:GVG:MAP-3xmCherry^35^. UBQ10p:mCitrine:1xPH^FAPP1^ and UBQ10p:mCitrine:P4M^SidM 26^, p35S ::RFP-HDEL^61^, pUBQ10p:DDRGK1:mCherry^62^. Lines SYT1-GFP/*sac7* and SYT1-GFP/mCherry-SAC7 were obtained by crossing. The following plasmids were previously published and characterized: UBQ10p:MAP:mCherry:*dm*OCRL and UBQ10p:MAP:mCherry:*dm*OCRL-dead^35^, UBQ10p:Lifeact:2xTU2^63^, UBQ10p:Lti6b-2xmCHERRY^64^, 35Sp:FKBP-mCherry:SAC1, 35Sp:FKBP:mCherry:SAC1dead and 35Sp:MAP:BFP2:FRB^42^, UBQ10p:mRuby:LiMETER^49^.

35Sp:SYT1-GFP/pGWB5, CFP-HDEL and 35S:GFP-NET3C plasmids were obtained from Miguel Botella (University of Malaga), Katharina Buerstenbinder (Leibniz Institute for Plant Biochemistry), and Steffen Vanneste (Ghent University), respectively.

Genotypes of individual plants were analyzed by PCR genotyping (for list of primers, see **Supplementary Data 1a**). Plants were grown in a growth chamber on peat-clay soil, watered with fertilizer as needed, under LED lighting (150 μmol.m-2.s-1) with a daily cycle of 16 hours of light (temperature 22°C and humidity 60%) and 8 hours of darkness (temperature 19°C and humidity 70%).

For *in vitro* culture conditions, seeds underwent surface sterilization by placing them in a sealed box with a solution composed of 4ml of 37% HCl in 100ml of bleach for a duration of four hours. Subsequently, the sterilized seeds were placed onto culture medium with 0.7% plant agar and 1% sucrose (pH 5.7), grown vertically in continuous light conditions at 21°C for 6 days before imaging. For *in vivo* root hair imaging and root hair phenotyping the seeds were placed onto a culture medium containing 2.15 g/L of MS (Duchefa, “0.5 MS”), 10 g/L of sucrose, 100 mg/L of myo-inositol, 500mg/L of MES hydrate, 0.5 mg/L of nicotinic acid, 0.5 mg/L of Vitamin B6, 0.1 mg/L of Vitamin B1, 2 mg/L of Glycine and 6 g/L Gelrite (Duchefa Biochemie) (pH 5.7). For *in vivo* observations of root hair growth, seeds were germinated in Lab-Tek II coverglass chambers (type of chamber:155360, Thermo Scientific) for 5 days.

### Plant transformation and selection

The constructs were transformed into C58 GV3101 *Agrobacterium tumefaciens* strain and selected on YEB media (5g/L beef extract; 1g/L yeast extract; 5g/L peptone; 5g/L sucrose; 15 g/L bactoagar; pH 7.2) supplemented with antibiotics. For transient transformation of *Nicotiana benthamiana* leaves, the Agrobacterium with the target construct was incubated at 28 °C for 48 h and then suspended in the infiltration media (10mM MgCl_2_ and 10mM TrisHCl pH=7.5). Then, the OD_600_ was adjusted to 1 using a spectrophotometer (Biophotometer, Eppendorf). For co-infiltration of several constructs, the same quantity of each transformed Agrobacterium was mixed to obtain a final OD_600_ of 1. Then the infiltration of the leaves was done as already described previously^65^.

For stable transformation of *Arabidopsis thaliana,* Agrobacterium cells with the target construct were solubilized in a solution containing 10mM MgCl_2_, 5% sucrose and 0.03% Silwet L-77 detergent. Flowering Arabidopsis plants were transformed using the floral dipping method^66^. The selection process was executed using the identification of red, green or blue fluorescent seeds (for plants carrying FastRed, FastGreen and Fast Cyan selection) or seedlings were grown on MS media with antibiotic or herbicide selection (50mg/L kanamycin, 30 mg/L hygromycin and 10 mg/L ammonium glufosinate).

### Cloning and generation of mutant lines

*Cloning of dmSAC1 constructs:* For cloning of UBQ10p:MAP:*dm*SAC1-mCherry construct, a synthetic gene (https://eu.idtdna.com/pages) corresponding to *dm*SAC1_1-521_ was codon optimized for expression in *A. thaliana* and flanked with BsaI recognition sites from both sides and linker sequence at the C terminus (for the full sequence, see **Supplementary Data 1b**). The gene was amplified and cloned into a pGGC000 Green Gate entry vector via BsaI restriction and ligation by Green Gate cloning strategy^67^. *dm*SAC1dead gene was designed by changing catalytic cysteine (388) to serine in the catalytic motif of the gene by PCR amplification of the sequences upstream and downstream of the catalytic cysteine with primer pairs whereof one primer has an altered nucleotide sequence at this site, followed by an overlap extension PCR to reconnect the gene fragments (for the list of primers, see **Supplementary Data 1a**). Then, *dm*SAC1dead was cloned into the GreenGate entry vector pGGC000 via BsaI restriction and ligation.

The UBQ10 promoter, the CDS of mCherry and Myristoylation and Palmitoylation sequence (MAP) were amplified by PCR using adequate primer pairs to add flanking BsaI restriction sites and matching overlaps for cloning into the GreenGate entry vectors pGGA000 (for UBQ10), pGGD000 (for mCherry), pGGB000 (for MAP), pGGE00 (for RBCS terminator) and pGGF000 (for Sulfadiazine resistance) via BsaI restriction and ligation.

For cloning of the UBQ10p:GVG:MAP:*dm*SAC1:mCherry construct, a UBQ10p:GVG (UBQ10pro:GVG-tE9::6xUAS-35Smini)^44^ construct (where GVG indicates the glucocorticoid receptor fused with the synthetic transcription activator VP16 and the GAL4 DNA binding domain) was cloned using Gibson cloning^68^ into pGGA000.The cloning of the rest of the construct was the same as described for UBQ10p: MAP-*dm*SAC1-mCherry above with a difference in pGGF000 vector that carried the Fast Cyan seed selection.

The expression cassettes were created with a GreenGate reaction using pSICE destination vector (carrying spectinomycin resistance) designed in our lab. All created entry vectors were confirmed by sequencing. For the full list of entry vectors and final constructs, see **Supplementary Data 2**.

*Generation of PI4Kα crispr mutant line:* Four CRISPR guides directed against PI4Kα were created using the website: http://crispr.hzau.edu.cn/CRISPR2/. For the guide sequences, see **Supplementary Data 1a**). The plasmid for transformation were prepared using the CRISPR-Cas9 cloning system and protocol developed in Stuttmann et al.^69^

### Imaging Configuration

Confocal imaging of five to seven days old Arabidopsis seedlings was conducted using a Zeiss microscope (vertical LSM 980 Axio Observer Z1/7 with AiryScan 2 module).

This setup utilized either 10X objective (numerical aperture 0.3) or 40x objective (numerical aperture 1.3, oil immersion) combined with GaAsP-PMT detector. For super-resolution imaging Airyscan2 module was applied using 40x objective.

The observations of transiently transformed leaves of *N. bethamiana* were all performed 2 days after infiltration, using a Zeiss confocal microscope (AxioObserver Z1) equipped with a spinning disk module (CSU-W1-T3, Yokogawa) and a Prime 95B camera (Photometrics). This setup utilized 40x objective (LD C-Apochromat 40×/1.1 W Corr M27). For live imaging of growing root hairs 100× Plan-Apochromat objective (numerical aperture of 1.46, oil immersion) was used.

Green Fluorescent Protein (GFP) was activated through a 488 nm laser, mCitrine marker was excited utilizing a 514nm laser, mCherry/RFP/mRuby marker was excited utilizing a 561nm laser, mTurquoise2/BFP2 marker was excited using 445nm laser. All emission wavelengths were detected between 300-735nm.

For optogenetic experiments, seedlings were grown in Lab-Tek II chambers for 5 days under continuous light. Before observation, the chambers were put in dark for 10 min. Then, the seedlings were observed using Zeiss confocal microscope (AxioObserver Z1, Carl Zeiss Group) equipped with a spinning disk module (CSU-W1-T3, Yokogawa) and a Prime 95B camera (Photometrics). This setup utilized 100x objective, and 4 minute-time lapses with 6-second interval between frames. The 561nm laser was applied during the whole-time lapse (for mRuby excitation), while the 488nm laser (70% power) was turned on after 2min (for the activation of the LOV2) and it was kept on until the end of the time lapse.

For Total Internal Reflection Fluorescence (TIRF) Microscopy, the inverted Zeiss microscope (AxioObserver Z1) was equipped with azimuthal TIRF iLas2 system (Roper Scientific) and a Prime 95 camera (Teledyne Vision Solutions). A 100x Plan-Apochromat objective (numerical aperture 1.46, oil immersion) was used. GFP was excited with a 488nm laser (150mW), and fluorescence emission was filtered by a 525/50 nm BrightLine® single-band bandpass filter. mCherry was excited with a 561 nm laser (80 mW) and fluorescence emission was filtered by a 609/54 nm BrightLine® single-band bandpass filter (Semrock,http://www.semrock.com/).

### Image analysis and quantification

Confocal images were analyzed using software ImageJ (https://fiji.sc/). For contact site density measurements, images were first segmented using ilastik machine learning-based software (https://www.ilastik.org/) where different pixel values were assigned to the ER, background and round structures (representing contact sites). For an example of input image and output segmented image see **Supplementary Fig.1**). Segmented images were then analyzed by ImageJ, using Fiji macro of our design (website link to the Fiji macro for contact site density quantification is listed in the **Supplementary Data 1c**).

*Quantification analysis of SYT1-SAC7 colocalisation*: Time lapse recordings were taken for a duration of 60 seconds at an interval of one frame every 1.3 seconds using TIRF microscopy. Time lapse images were processed in FIJI software using a custom macro script as following: SYT1-GFP contact sites were segmented in the first time point using a differencial of Gaussian filtering. Kymographs were then obtained for each contact site (line length 3.3µm, 0.55µm line width), before being concatenated into a single image using the “Combine” function. In a second step, colocalisation was quantified using a Fiji macro of our design. SUM-projection were applied to differencial of Gaussian filtered time-lapse acquisitions and then subsequently analyzed using JACOP plugin (https://imagej.net/plugins/jacop) to calculate Pearson ‘s coefficient and DiANA signal overlapping. (https://imagej.net/plugins/distance-analysis). For an example of representative input images, see **Supplementary Figs.5c-d**. For website links of Fiji macros used in this study, see **Supplementary Data 1c**.

*Root hair quantification analysis:* Root hair length and density was assessed in 5 days old seedlings. Root hairs falling within the focal plane, spanning from 2000μm to 4000μm distance from the root apex, were subjected to measurement. Root hair density and length were analyzed in ImageJ using Fiji macro of our design (website link to the Fiji macro for quantification of root hair density and length is listed in the **Supplementary Data 1c**).

*Quantification of cortical index*: Cortical index was calculated manually with a draw-by-hand fluorescence extraction ROI method on Fiji. The values of the ROI fluorescence (apical, side, perinuclear) was subtracted by the value of the background of the same cell. For each cell, the cortical index is calculated as the mean apico-side fluorescence divided by the perinuclear fluorescence. For more details, see **Supplemental Data1d**.

### Statistical analysis

All statistical analyses were done using R (version 2024.12.0+467) and the RStudio interface.

### Co-Immunoprecipitation and Immunoblotting

The experiment was performed on 8 days old transgenic seedlings plants. All seedlings were frozen using liquid nitrogen and ground into a fine powder. 500 mg of ground tissue was used for each sample. The proteins were extracted with the extraction buffer at 2mL/g of powder (50 mM Tris–HCl, pH 7.5; 150 mM NaCl; 10% glycerol; 10 mM EDTA, pH 8; 1 mM NaF; 1 mM Na_2_MoO_4_.2H_2_O; 10 mM DTT; 0.5 mM PMSF; 1% (v/v) P9599 protease inhibitor cocktail; Nonidet P-40 0.5% (v/v.)). The suspension was mixed using a tilting rocker during 30min at 4°C. The solution was then centrifuged for a duration of 30min at 4°C at 16000g. The resulting supernatants were filtered with Poly-prep chromatography columns (BioRad^TM^) where 100 µL was kept for the input of the immunoblot. The other supernatants were diluted in the extraction buffer without Nonidet P-40 at 1:1. The resulting concentration of Nonidet P40 was 0,25%. The protein samples were subsequently incubated for 2 hours at 4°C on a tilting rocker with 20µL ChromoTek^TM^ RFP-Trap Magnetic Agarose beads. The beads were collected and washed with the IP extraction buffer with 0.05% Nonidet P-40, four times. After washing, the beads were resuspended in 75µL of 2x concentrated Laemmli buffer (Sigma-Aldrich^TM^) and incubated for 45 minutes at 70°C. Finally, 20µL of the Input total, IP and Co-IP proteins were loaded into a gradient 4-20% MP TGX Stain-Free SDS-Page acrylamide gel (BioRad^TM^) and the proteins were separated by eletrophoresis. For immunoblotting, the proteins separated by the Acrylamide gel electrophoresis were transferred onto a Low Fluorescence PVDF membrane (Immun-Blot^®^, BioRad^TM^) using the Trans-Blot Turbo RTA transfer kit (BioRad^TM^). The protocol used for the transfer was the manufacturer recommendation for mixed molecular weight proteins (Trans-Blot Turbo, BioRad^TM^). Subsequently, the protein-containing PVDF membranes were incubated with rabbit polyclonal anti-SYT1 antibody (1:5000) for the detection of Co-IP SYT1 epitope, or Chromotek^TM^ mouse anti-RFP monoclonal IgG2c antibody (clone 6G6, 1:2000) for the detection of mCherry epitope.

The secondary antibody used, allowing the visualization of the presence of SYT1 or mCherry proteins on the PVDF membrane, was an anti-rabbit IgG whole molecule-Peroxidase (1:80,000; A0545, Sigma-Aldrich^TM^) or anti-mouse IgG whole molecule-Peroxidase (1:80,000; A9044, Sigma-Aldrich^TM^), respectively.

Proteins and tagged proteins on the immunoblot were visualized using the Clarity ECL Western Blotting Substrate (BioRad^TM^) or SuperSignal West Atto (Thermo Fisher Scientific^TM^). The images of the immunoblot were acquired with the Chemidoc MP System (BioRad^TM^). The stain-free imaging (BioRad) of the PVDF membrane blot was used to confirm equal loading of the different samples.

## Acknowledgments

We thank the SiCE group for discussion and Claire Lionnet for help with microscopy. We would like to thank Katharina Buerstenbinder and Steffen Vanneste for sharing their CFP-HDEL and 35S:GFP-NET3C plasmids, respectively. We are grateful to Pengwei Wang for the UBQ10p:mRuby:LiMETER line. We thank Y.Boutté (LBM, Bordeaux), A. Martiniere (IPSiM, Montpellier) and J. Pérez-Sancho (University of Málaga) for the insightful feedback on the manuscript.

## Author contributions

Conceptualization: Y.J. and V.M. Methodology: V.M., V.B., G.D., F.R., V.A.S., M.A.B., and Y.J. Investigation: V.M., G.D., V.B., F.R., V.A.S., J.M.L., S.G., and S.G.H. Funding acquisition: Y.J., V.M., V.A.S. and M.A.B. Visualization: V.M. Writing: V.M and Y.J. Supervision: Y.J. and M.A.B.

