## Supplementary Data 1 for "Feedback between PI4P signaling and ER-PM contact sites orchestrates polarized root hair growth"

### Supplementary Data 1a. List of primers used in this study.

| ***Plant genotyping*** | |
|---|---|
| sac7-3 LP | TTGTTCCAAAGAAACCTCGG |
| sac7-3 RP | GAGACGGTTGATTCTCGGAAC |
| syt1-2 LP | AGGTCTCGCGATTTATTAGGG |
| syt1-2 RP | GCCTCCTGACAAGTATAGGGG |
| syt3 LP | ATCCTCTGGTGGTTTCAAAGG |
| syt3 RP | GGTTGATCTCAGGTATGTGCC |
| Salk Lbb1.3 | ATTTTGCCGATTTCGGAAC |
| Sail Lbb1.3 | TAGCATCTGAATTTCATAACCAATCTCGATACAC |

| ***Cloning of Drosophila SAC1dead constructs*** | | |
|---|---|---|
| SAC1dead_part1 FW_C | aaaaGGTCTCaGGCTctATGGATAGCCGTGAGGAGAA | dmSAC1 dead into pGGC000 |
| SAC1dead_part1 RV_C | tctcGGTCTCaTGAAAACGCCTGTTTGCGTTGATA | dmSAC1 dead into pGGC000 |
| SAC1dead_part2 FW_C | tctcGGTCTCaTTCAGGACCAATtctATCGACTGT | dmSAC1 dead into pGGC000 |
| SAC1dead_part2 RV_C | ttttGGTCTCaCTGAACGTCCAGCTCCTCCAG | dmSAC1 dead into pGGC000 |
| UBQ10_GVG_fw_A | tctcGGTCTCaACCTttAGTCTAGCTCAACAGAGCTT | UBQ10prom+GVG into pGGA000 |
| UBQ10_GVG_rv_A | tctcGGTCTCaTGTTctgctttttgtacaaacttgc | UBQ10prom+GVG into pGGA000 |
| mCherry_stop_FW_D | tctcGGTCTCaTCAGAAATGGTGAGCAAGGGCGAGgag | mCherry+stop codon into pGGD000 |
| mCherry_stop_RV_D | tctcGGTCTCaGCAGCTACTTGTACAGCTCGTCCATG | mCherry+stop codon into pGGD000 |

| ***The guide sequences for pi4kα crispr mutant design*** | |
|---|---|
| Guide 1 (79 bases de l'ATG) | GTCGTTGTCCACAAACGGAA |
| Guide 2 (314 bases de l'ATG) | GAGTTATCTTTCGTGTGCGG |
| Guide 3 (629 bases de l'ATG) | TAGTACGAACCGTTATCCCTC |
| Guide 4 (1493 bases de l'ATG) | GGTACTAGCTGTCTGTGCAC |

| ***Cloning of the 35S:GFP-NET3C (Steffen Vanneste, Ghent University)*** | |
|---|---|
| NET3C_FW | GGGGACAAGTTTGTACAAAAAAGCAGGCTTCCCGCCAATGGTTAGAGAAGAGGAGAAATC |
| NET3C_REV | GGGGACCACTTTGTACAAGAAAGCTGGGTCCTAAAGGACCTTGTTGCCATC |

### Supplementary Data 1b. The sequence of codon optimized *Drosophila* SAC1$_{1\text{-}521}$ gene.

*dm*SAC1$_{1\text{-}521}$

ATGGATAGCCGTGAGGAGAACGCGGTTTATGACGACATGAACTTGTACATCGCTCCACAGAGTTTCATAATCGAGCCCAATGGTGGCGACGAGCT
CCTCGTTATAGGAAGGCATGATAAGGTAACCAGAGTACAGCCGGCGAGCGGGGGCCTTGTTGCTAACCTAAGGCCAACACGTCGAATCTGTGGC
GTTCTCGGAACCATTCACTTGTTATCATGCGATTATCTGTTAGTAGCCACTCACCGTTTATTTGTAGGTGTGCTAAATGGGGCCGTTGTCTGGCGTC
TCGCGGGATACGACATTATACCTTACATTCCCAATTCCTTCCAGCGTAAGGAGAACGAGAACTACCTAAGGCTTCTACGACAGACTCTCGACACCA
AATTCTTCTATTTTAGTTACAGATATGATCTTACCAACTCTTTACAACGTCAGAGGGAAGTTGCACAGTCCAGACCCGAAGTATCAGGCCTTCTTCAG
AGGGCCGAACAACGTTTTGTATGGAATGGCTATGTCCTTAGGCAGTTTAACTGTGATAAGATGGAGAAGTTTCAGCTTCCGCTGGTTCTAGGATTT
GTTAGCATCAATCAGGTTCAGATAAATGGCCAGACCTTTTTTTGGAGTATCATAACGCGTCGAAGCGTACAGAGGGCTGGAACCCGTCTGTTCTGC
CGTGGAAGTGATGAGCAAGGGCATGTCGCAAACTTTGTAGAaaCCGAGCAAATTGTGGAATTCAACGGGCAACTTACGGGATTCGTACAAACACGT
GGTAGCATGCCTTTTCACTGGCATCAACTCCCTAATTTGCGTTACAAACCAAGGCCGGTGTTAGTTCCGGGAAAGGATCACCTGGCAGCTTGTGGa
CTTCATTTCAAAGAGCAGATTAGATTATATGGAAACAACGTTGCGGTCAACCTCGTTGATCACAAAGGCGCTGAAGGCGAGCTTGAAGCCACGTAT
GCGCGTCTCGTTCGAGAAATGGGCAATCCACAGGTGCGATATGAGTCATTCGATTTTCACAGCGAGTGCAGAAAGATGCGTTGGGACAGACTTAA
CATTTTGATAGACCGATTAGCCCACGAACAAGATCAATTTGGGGTTTATCATGTCTTTGACGACGGCAAGCTGGTATCAACGCAAACAGGCGTTTTC
AGGACCAATTGCATCGACTGTCTTGACAGAACAAATGTCGTACAGTCCATGTTAGCGAGACGAAGTCTCACAGCTGTCCTTCAGAAATTAGGGGTC
CTCCATGTCGGGCAAAAGGTAGAGCATGCAAGCGATATCTTCGAAAGCATCTTTAAAGGCGTTTGGGCTGACAATGCCGACTTAGTGAGCTTGCAA
TACTCCGGAACATGTGCTTTGAAAACCGATTTTACCCGTACTGGCAAGAGGACAAAGAGTGGCGCCATGCAAGATGGCAAGAACTCCCTTATGAGA
TACTACCTAAACAACTTCGCCGATGGACAGAGACAGGACTCAATTGATCTGTTCCTTGGAAAATACTTGGTAAATGACAATGAAGGCGGTGCAGTC
CCCAGCCCGCTAGAATCTAAACACGGGGGAGGTACTGCTCGTGGAGCTGCTGCTGGAGCTGGAGGAGCTGGACGT

Red letters represent the catalytic motif and the sequence encoding for the catalytic cysteine is highlighted in yellow. For designing the *dm*SAC1dead, at the place of catalytic cysteine, serine was introduced by changing TGC sequence to TCT sequence. Blue letters indicate the linker sequence.

### Supplementary Data 1c. Website links to Fiji macros used in this study.

| | |
|---|---|
| Fiji macro for contact site density quantification | https://github.com/RDP-vbayle/SiCE_FIJI_Macro/blob/main/misc/MAcroPatternERcs%20Markovic%20et%20al.%202026.ijm |
| Fiji macro for quantification of the root hair length and density | https://github.com/RDP-vbayle/SiCE_FIJI_Macro/blob/main/misc/MacroRootHair%20Markovic%20et%20al.%202026.ijm |
| Fiji macro for contact site segmentation of SYT1-GFP-mCherry-SAC7 time-lapses | https://github.com/RDP-vbayle/SiCE_FIJI_Macro/blob/main/misc/MAcroSAC7SYT1Kymo%20Markovic%20et%20al%202026.ijm |
| Fiji macro for colocalization analysis of SYT1-GFP and mCherry-SAC7 | https://github.com/RDP-vbayle/SiCE_FIJI_Macro/blob/main/misc/MAcroSAC7SYT1coloc%20Markovic%20et%20al%202026.ijm |

**Supplementary Data 1d. Quantification of the cortical index.**

An example of a root epidermal cell:

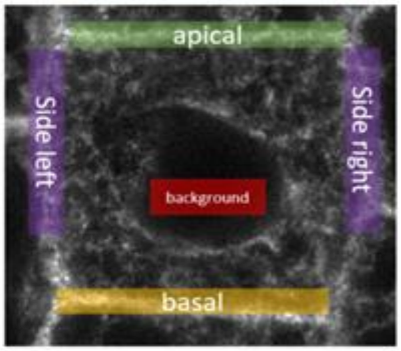


**Cortical index:**

$$\frac{(\text{Apical Fluo} - \text{Background}) + (\text{Side right} - \text{Background})}{2x\ (\text{Perinuclear Fluo} - \text{Background})}$$
