## Supplementary Information for "Feedback between PI4P signaling and ER-PM contact sites orchestrates polarized root hair growth"

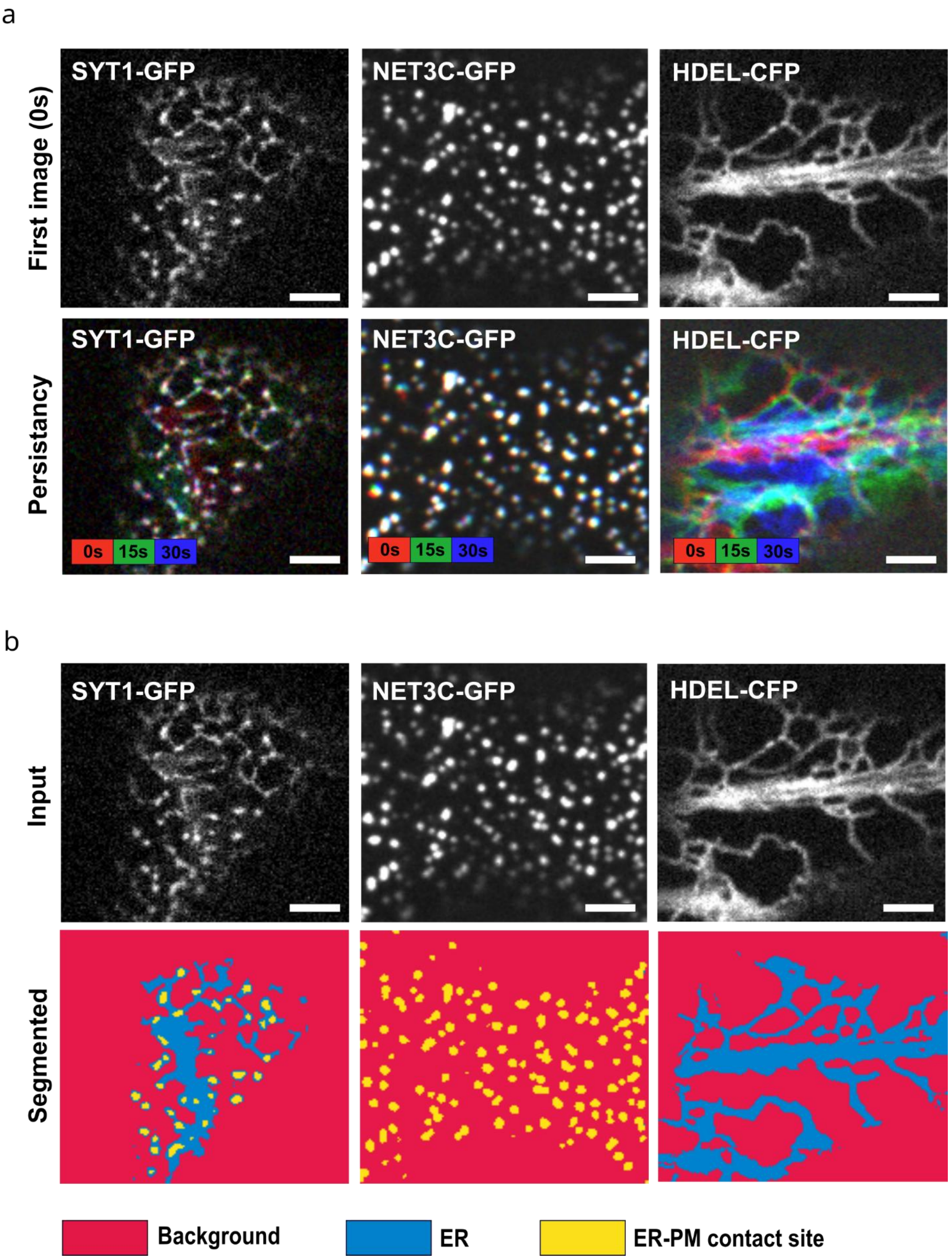
a
SYT1-GFP
NET3C-GFP
HDEL-CFP
First image (0s)
Persistancy
0s
15s
30s
b
Input
Segmented
Background
ER
ER-PM contact site

**Supplementary Figure 1.** SYT1-GFP and NET3C-GFP localise in stable ER-PM contact sites that can be segmented by the ilastik software **(a)** Upper panels: Cortical view of *N.benthamiana* leaf epidermal cell expressing SYT1-GFP (left), NET3C-GFP (middle), and HDEL-CFP (right) at 0 s of a 30-s time-lapse series. Lower panels: Merged pseudo coloured images of SYT1-GFP, NET3C-GFP, and HDEL-CFP at 0, 15, and 30 s, showing signal persistence over time (persistency mapping). Unlike the dynamic ER marker HDEL-CFP, SYT1-GFP and NET3C punctate structures remained stable throughout the time course. Thirty frames were acquired at 1-s intervals. **(b)** Segmentation of stable SYT1-GFP and NET3C round structures using the machine-learning software ilastik. Upper panels: Input images of SYT1-GFP, NET3C-GFP, and HDEL-CFP**.** Lower panels**:** ilastik-segmented images. The software successfully distinguished ER structures (blue), stable punctate/round structures (yellow), and background signal (red). Scale bars = 5µm.

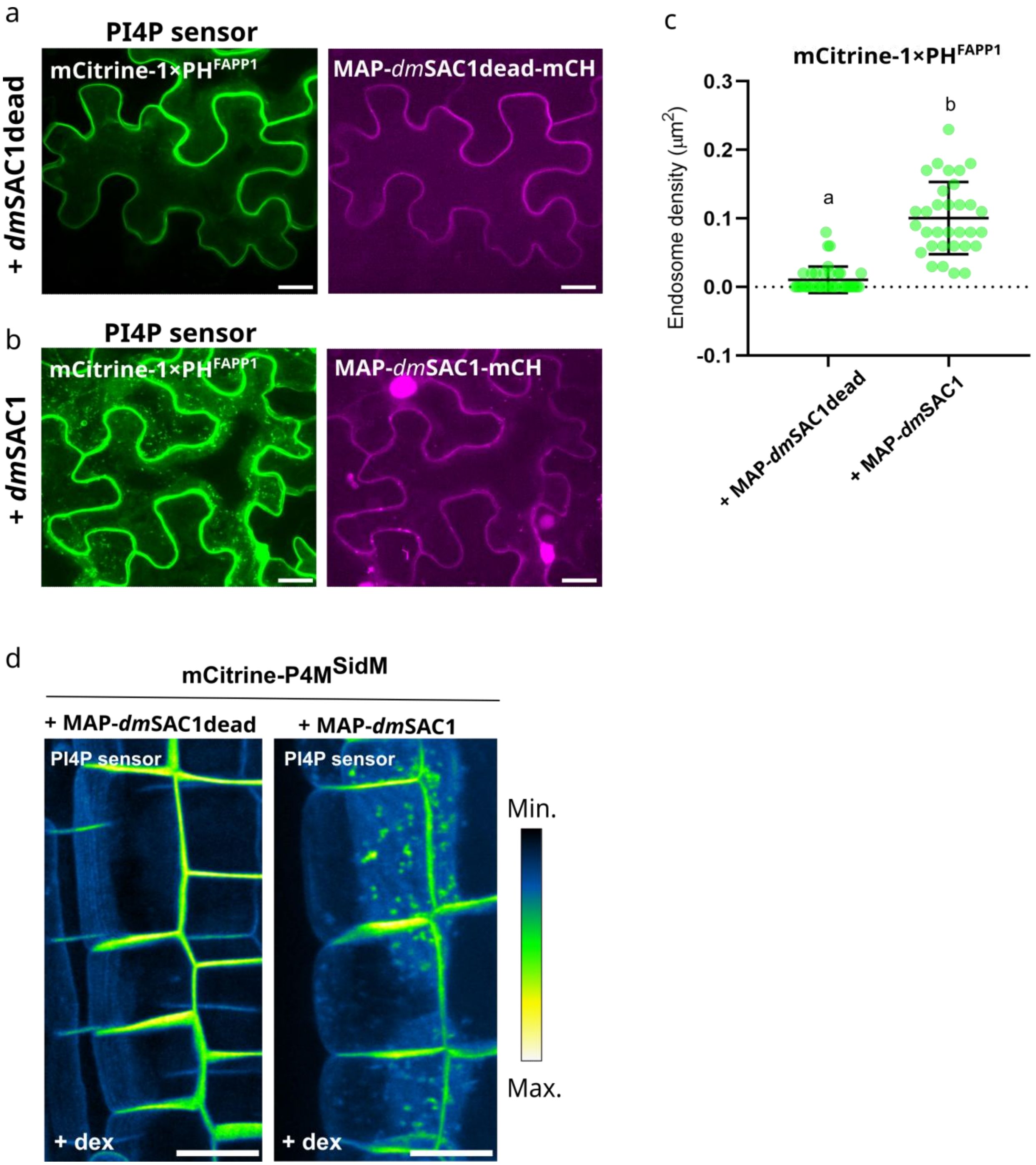


**Supplementary Figure 2.** MAP-*dm*SAC1-mCherry efficiently depletes the PI4P in *Nicotiana benthamiana* leaves and *Arabidopsis thaliana* roots. **(a)** Representative confocal images (maximum intensity projection) of the *N. benthamiana* leaf epidermal cell expressing MAP-*dm*SAC1dead-mCherry (right panel) and PI4P sensor, mCitrine-1x

PH$^{FAPP1}$ (left panel). **(b)** Representative confocal images (maximum intensity projection) of the *N. benthamiana* leaf epidermal cell expressing MAP-*dm*SAC1-mCherry (right panel) and PI4P sensor, mCitrine-1x PH $^{FAPP1}$ (left panel). **(c)** Quantification of endosome density of **(a)** and **(b).** Bold, horizontal lines represent the means, whereas error bars represent SD. Letters denote statistically different groups calculated by one-way ANOVA with post hoc Tukey's honest significant difference test; $p < 0.01$. At least sixteen cells ( n>16) per group were used for quantification. **(d)** Maximum intensity projection of *Arabidopsis* root cells expressing a PI4P biosensor mCitrine-P4M$^{SidM}$ and MAP-*dm*SAC1dead (left panel) and MAP-*dm*SAC1 (right panel) 16h of dex induction. Scale bars= 20µm.

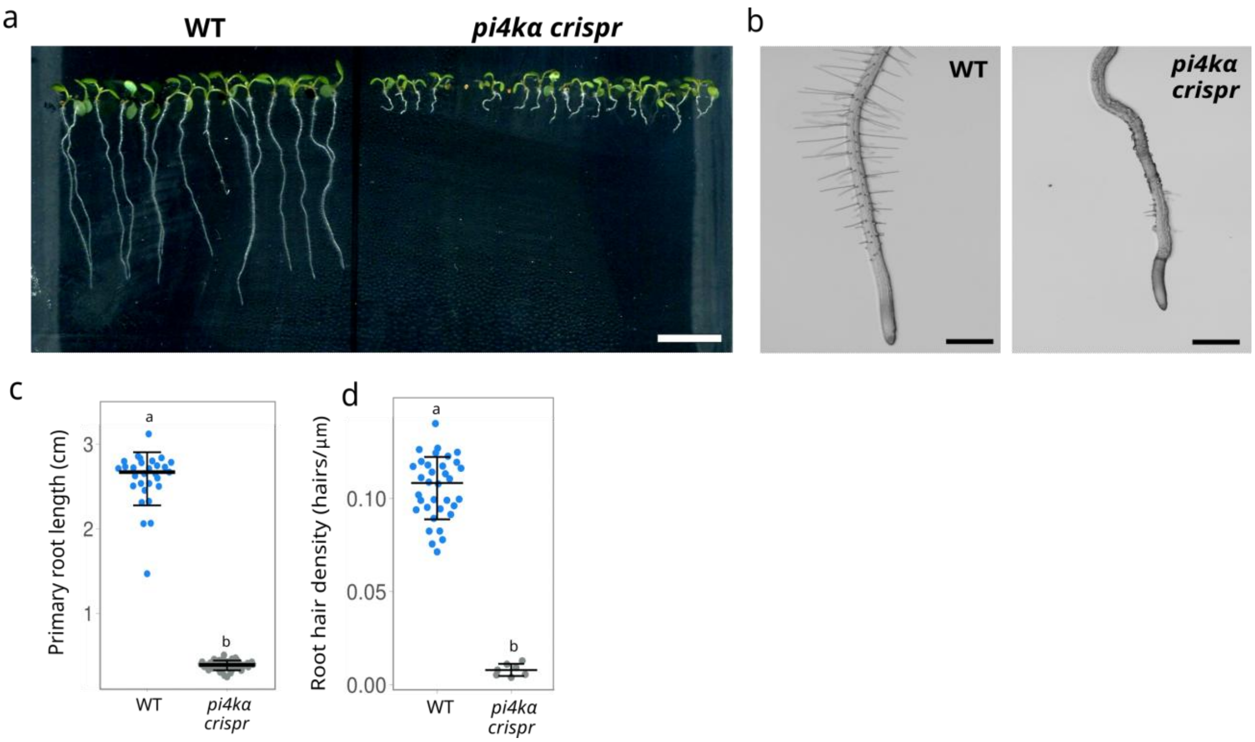


**Supplementary Figure 3.** Tissue-specific *pi4kα crispr* plants exhibit defects in root and root hair development. **(a)** Representative image of a plate with 10-days old wt (left) and *pi4kα crispr* (right) Arabidopsis seedlings. Scale bar = 1cm. **(b)** Representative images of wt (left) and *pi4kα crispr* (right) Arabidopsis roots. Scale bars = 500μm. **c-d** Quantification of the **(c)** primary root length and **(d)** root hair density of seedlings presented in **(a)** and **(b),** respectively. Letters denote statistically different groups calculated by one-way ANOVA with post hoc Tukey's honest significant difference test, $p < 0.01$. Horizontal lines represent the medians, whereas error bars represent SD. Each dot represents an individual **(c)** root or **(d)** root hair, and n refers to the total number of analysed roots and root hairs.

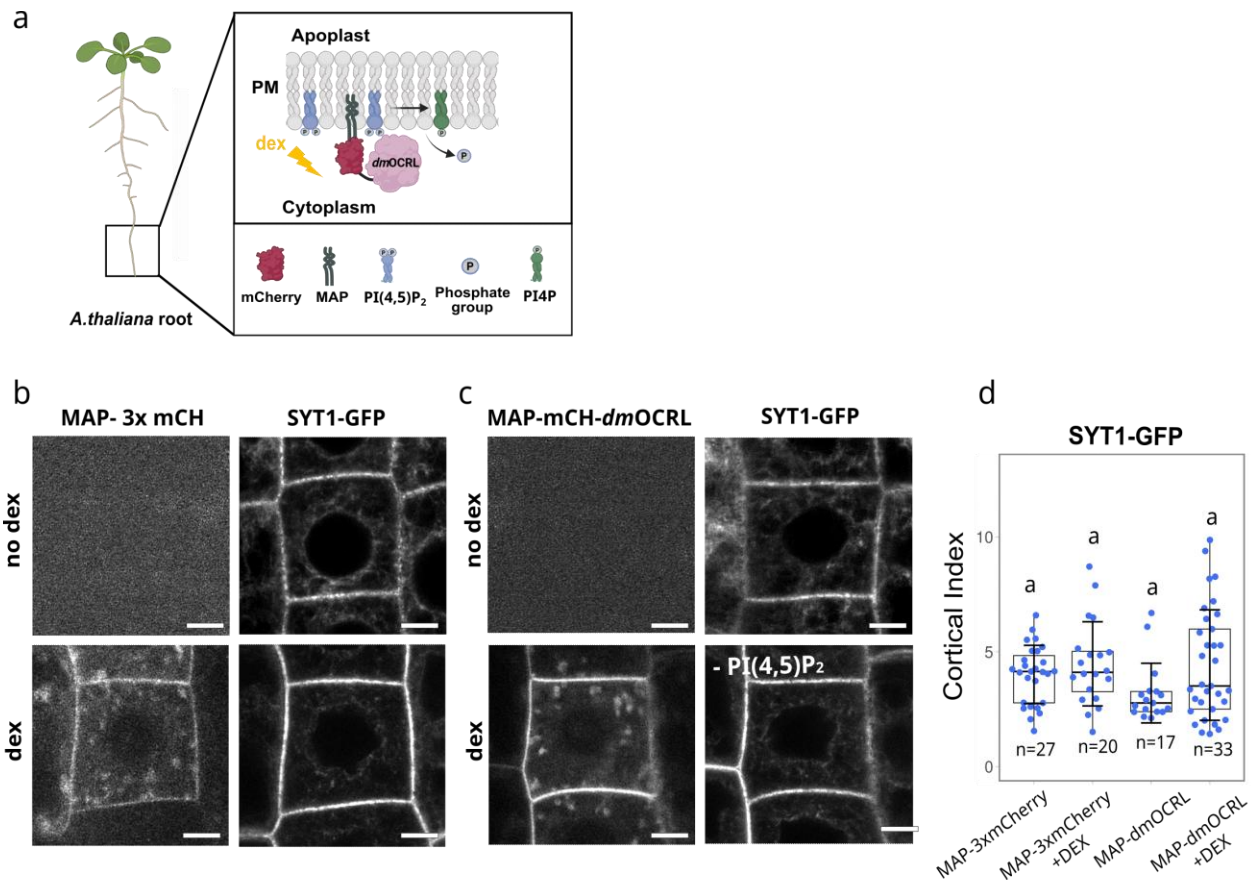


**Supplementary Figure 4.** $PI(4,5)P_2$ is dispensable for establishment of SYT1-mediated

ER-PM contact sites in Arabidopsis root cells **(a)** Schematic representation of the dexamethasone (dex)-inducible iDePP system for plasma membrane PI(4,5)$P_2$ depletion by *dm*OCRL in Arabidopsis cells. **(b)** Representative images of the fluorescent signal corresponding to MAP-3x-mCherry (left upper panel) and SYT1p:SYT1:GFP (right upper panel) in the same cell, without dex treatment, and MAP-3x-mCherry (left down panel) and SYT1p:SYT1:GFP (right down panel) in the same cell after 16 hours of dex treatment. **(c)** Representative images of the fluorescent signal corresponding to MAP-mCH-*dm*OCRL (left upper panel) and SYT1p:SYT1:GFP (right upper panel) in the same cell, without dex treatment, and MAP-mCH-*dm*OCRL (left down panel) and SYT1p:SYT1:GFP (right down panel) in the same cell after 16 hours of dex treatment. **(d)** Statistical analysis of cortical index of **(b)** and **(c)**. Horizontal lines represent the medians, whereas error bars represent SD. Each dot represents an individual cell, and n refers to the total number of analysed cells. Different letters indicate statistically significant differences between groups calculated by Kruskal–Wallis test. Scale bars = 5 µm.

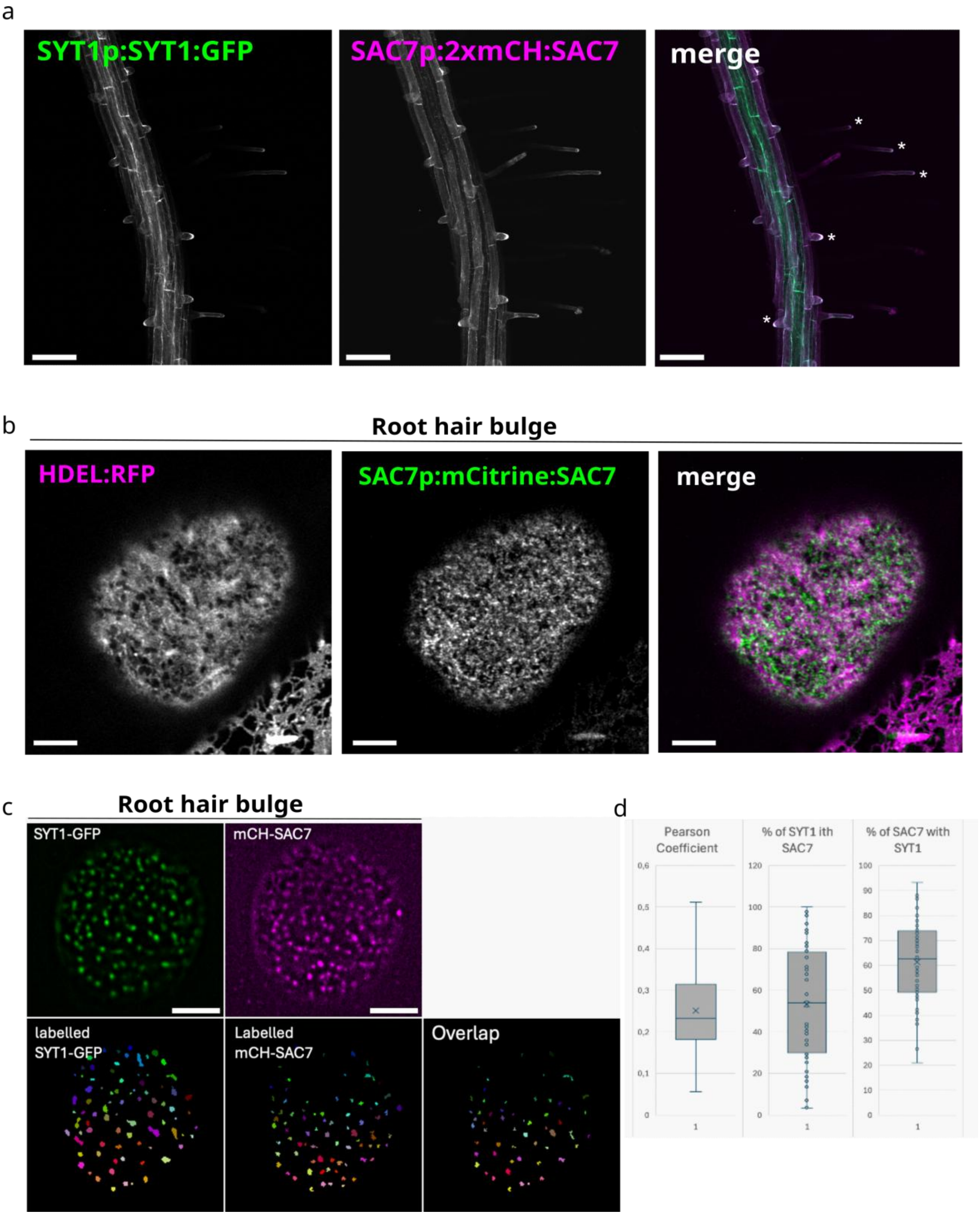


**Supplementary Figure 5.** SAC7 associates with SYT1 in root hair bulge and tip of the growing root hairs. **(a)** Representative image of 7 days old Arabidopsis root expressing SYT1-GFP and SAC7-mCherry. White asterisks indicate colocalization of SYT1-GFP and SAC7-mCherry in root hair bulge and at the tip of growing root hairs. Scale bars = 100 µm. **(b)** Cortical view of a root hair bulge expressing ER bulk marker HDEL-RFP and mCitrine-SAC7. Scale bars= 5 µm. **(c)** Upper panels: SUM- projection of time lapses acquired with TIRF microscopy (67 frames-total time 40s) Scale bars = 5 µm. Down panels: Round objects presented in upper panels were segmented and SYT1-GFP and mCherry-SAC7 overlapping objects were examined using ImageJ DiANA plugin. **(d)** Quantification of co localization and overlapping SYT1-GFP and mCherry-SAC7 objects using ImageJ DiANA plugin (n=61 root hair bulges).

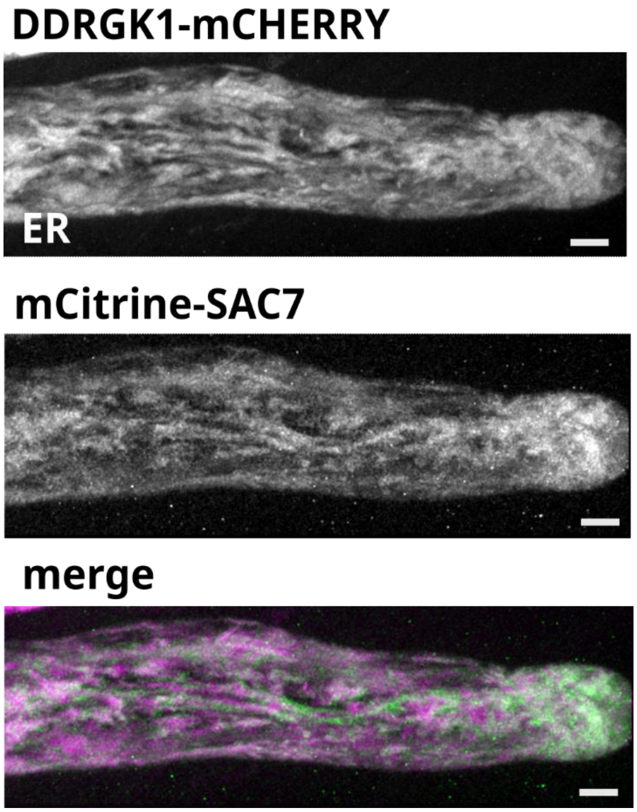


**Supplementary Figure 6.** The ER membrane marker and SAC7 accumulate in growing root hairs. Maximum intensity projection of a root hair expressing the DDRGK1-mCherry ER membrane marker and SAC7p:mCitrine:SAC7. Scale bars = 10 µm.

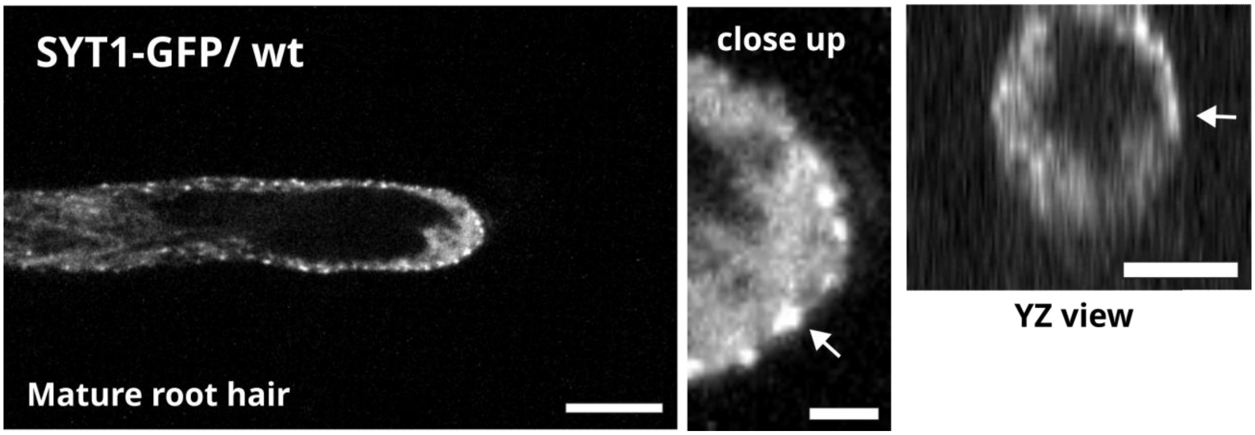


**Supplementary Figure 7.** SYT1 contact sites localise at the tip in the mature root hairs. Maximum intensity projection of a mature (non-growing) root hair expressing SYT1-GFP.  Scale bars = 10 µm. The arrows on the close up and YZ view images indicate accumulation of SYT1-GFP at the root hair tip. Scale bars=5 µm.

**Supplementary Videos**

**Supplementary Video 1.** Time lapse of a root hair expressing SYT1-GFP in wild-type background, related to **Fig.7a**. Images were taken every 10s. Note that SYT-GFP localizes in the ER at the tip of the growing root hair, with no presence of contact sites.

**Supplementary Video 2.** Time lapse of a root hair expressing SYT1-GFP in *sac7* background, related to **Fig.7b**. Images were taken every 10s. The presence of SYT1-GFP contact sites at the tip of growing root hair is indicated by the yellow arrow.

**Supplementary Video 3.** Time lapse of a mature (non-growing) root hair expressing SYT1-GFP in wild-type background, related to **Supplementary Fig.7**. Images were taken every 10s. The presence of SYT1-GFP contact sites at the tip of non-growing root hair is indicated by the yellow arrow.

**Supplementary Video 4.** Time lapse of a cortical region of root epidermal cell expressing mRuby-LiMETER. Images were taken every 3s. After blue light activation, mRuby-LiMETER localized in stable puncta representing ER-PM contact sites.

**Supplementary Video 5.** Time lapse of a root hair expressing mRuby-LiMETER, related to **Fig.7f**. Images were taken every 6s. Blue light was applied at 120s time point.

**Supplementary Video 6.** Time lapse of a root hair expressing Lti6b-mCherry related to **Fig.7g**. Images were taken every 6s. Blue light was applied at 120s time point.
